# An *in vitro, in utero* and *in silico* framework of oxygen diffusion in intricate vascular networks of the placenta

**DOI:** 10.1101/2021.12.01.470714

**Authors:** Nikhilesh Bappoo, Lachlan J. Kelsey, Yutthapong Tongpob, Kirk W. Feindel, Harrison Caddy, Caitlin S. Wyrwoll, Barry J. Doyle

## Abstract

The placenta is a temporary and complex organ critical for fetal development through its subtle but convoluted harmonization of endocrine, vascular, haemodynamic and exchange adaptations. Yet, due to experimental, technological and ethical constraints, this unique organ remains poorly understood. *In silico* tools are emerging as a powerful means to overcome these challenges and have the potential to actualize novel breakthroughs. Here, we present an interdisciplinary framework combining *in vitro* experiments used to develop an elegant and scalable *in silico* model of oxygen diffusion. We then use *in utero* imaging of placental perfusion and oxygenation in both control and growth-restricted rodent placentas for validation of our *in silico* model. Our framework revealed the structure-function relationship in the feto-placental vasculature; oxygen diffusion is impaired in growth-restricted placentas, due to the diminished arborization of growth-restricted feto-placental vasculature and the lack of decelerated flow for adequate oxygen diffusion and exchange. We highlight the mechanisms of impairment in a rat model of growth restriction, underpinned by placental vascular impairment. Our framework reports and validates the prediction of blood flow deceleration impairment in growth restricted placentas with the placenta’s oxygen transfer capability being significantly impaired, both globally and locally.

## Main

Computational experimentation is an emerging tool that is increasingly being leveraged to elucidate complex structure-function relationships in a number of biological systems such as the lungs ^1, 2^, placenta ^3-5^, tumors ^6, 7^ and others ^8-11^. This *in silico* approach overcomes challenges of conventional experiments, allowing for novel measurements to be made or experimental interventions to be actioned, both of which would otherwise be difficult, if not impossible to achieve conventionally. Contributing to this advancement are novel high-resolution imaging approaches, both *in vivo* and *ex vivo*, which have informed the development of more realistic modelling frameworks which are however often limited by the lack of validation data.

In obstetrics, fetal growth restriction is a poorly diagnosed and understood pathology driven by gas exchange dysfunction in the placenta. This unique organ develops an expansive vascular network of over 500km in length during pregnancy ^12^ to mediate efficient oxygen, nutrient and waste transport at the fetal-maternal interface and hence drive growth and health outcomes of the developing fetus. It has been hypothesized to be strongly influenced by synergistic structural, haemodynamic and diffusion impairment ^3-5, 13^, however, ethical constraints of pregnancy research and the low resolution of routine ultrasound limit the extent of emerging research into the human placenta.

Consequently, animal models, particularly rodents which share similarities to human placenta ^14^, offer an avenue to provide unique insights into both health and disease ^15-17^. Seminal studies have included *ex vivo* casting and imaging experiments, ^18-21^, *in silico* flow or oxygen diffusion simulations ^18, 22-26^ and more recently, *in utero* placental magnetic resonance imaging (MRI) ^27-29^. Nonetheless, this research is vastly limited by the complexity of placental vasculature and the difficulties in experimental validation of computational tools, which make it challenging to quantify and validate the local variability in perfusion and oxygenation in both *in utero* and *in silico* approaches ^30^. Although these novel methods hold immense potential to shed light on the key mechanics of placental gas exchange and impairment, they are rarely used in tandem to overcome current limitations and generate novel, more sensitive and more specific indicators.

In this study, we present an elegant and scalable computational framework (Figure 1) applicable to realistic geometries, incorporating the biological complexities of gas diffusion and accounting for experimental data for both model verification and validation. We present an application in a rat model of late gestation placentae in control and growth restricted contexts. We use *in utero* flow measurements from previous work ^13^ to model flow in subject specific placental arteries (collected *ex vivo*), a combination of *in vitro* and *in silico* experiments to develop verified oxygen diffusion simulations in bioinspired capillaries, and *in utero* placental MRI to validate *in silico* findings. We demonstrate that through extensive verification and validation using a combination of *in vitro, in utero* and *in silico* methods, we can generate novel insights on poorly understood structure-function relationships in vascular systems such as the placenta, and that both structure and haemodynamics influence diffusion impairment in disease.

**Figure 1:**
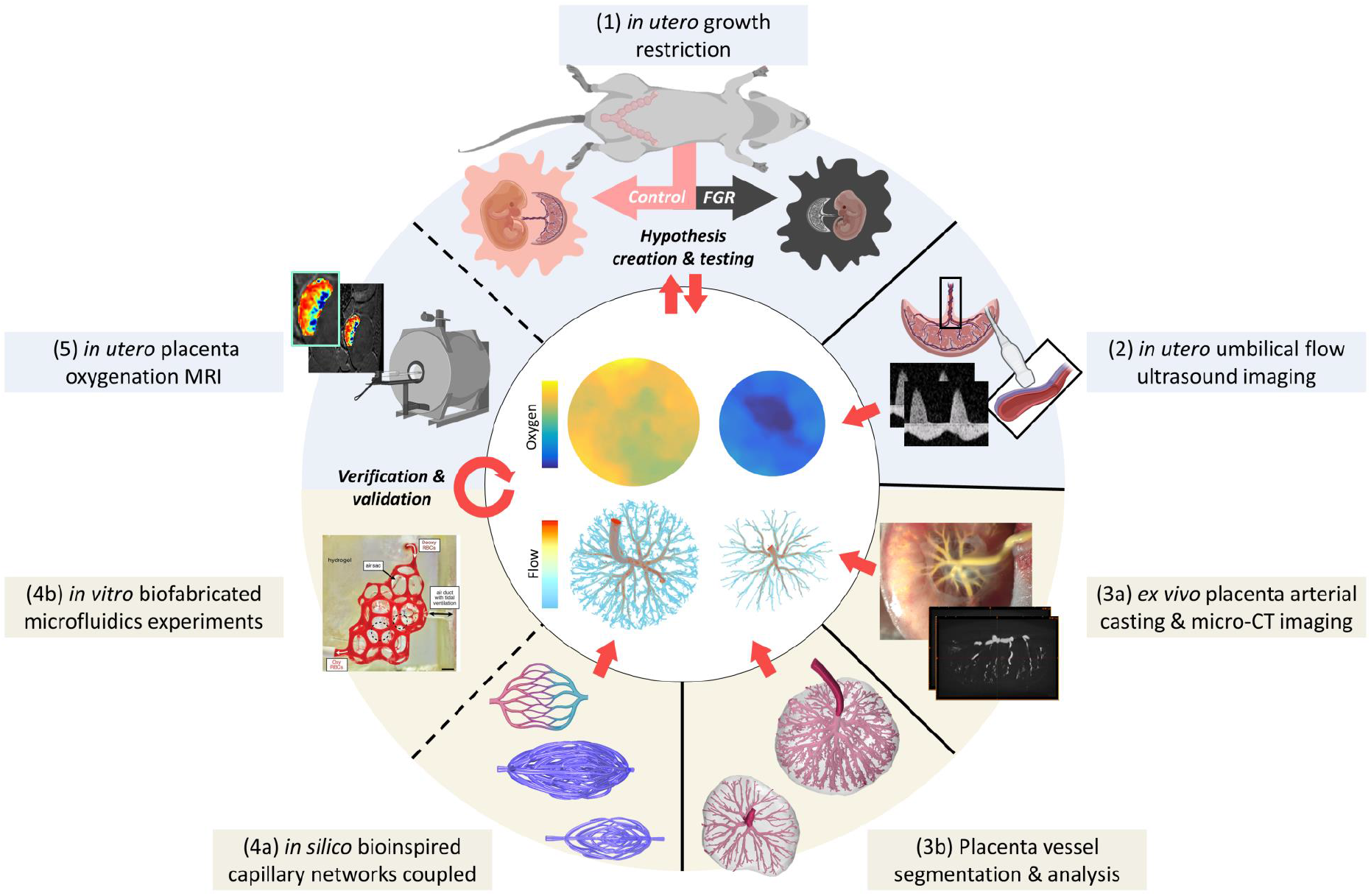
Pipeline for *in vitro, in utero* and *in silico* research into 3D placental structure, haemodynamics and function. A well-established glucocorticoid (dexamethasone or dex exposure) model of fetal growth restriction (FGR) (1) in rats is used to investigate mechanisms of placental impairment during late pregnancy compared to control rat pregnancies. Umbilical artery flow measurements (from Bappoo et al.^13^) are taken *in utero* using High Frequency Doppler Ultrasound imaging (2) to define model inputs for *in silico* computational fluid dynamics (CFD) simulations. Feto-placental units are resected and blood is cleared before the placental arterial tree is cast using a radiopaque casting agent and subsequently imaged using micro-CT (3a). The placental tissue is reconstructed in 3D (specifically, arteries) and vessel structure is analyzed using a 3D rooted tree approach to extract geometrical data such as vessel size (3b) ^13, 20^. Openly-sourced bioinspired capillary networks ^31^ are constructed *in silico* (4a) and oxygen diffusion is characterized for the capillary geometries before coupling with terminal vessel outlets. Bioinspired vascular networks are fabricated using hydrogels and microfluidics experiments are performed ^32^ to measure oxygen gained for specified flow rates, hence ensuring that *in silico* oxygen diffusion modelling methods are verified (4b). The 3D geometries are prepared for flow simulations to compute the steady-state blood flow and oxygen diffusion at the fetal-maternal interface (center figure) and placental function can be compared between experimental groups. Using MRI, ΔT2* values are calculated to provide a relative measure of overall oxygenation *in utero* and is compared to *in silico* results for validation (5). Here, this framework is used to elucidate structural, haemodynamic and functional (oxygen diffusion) changes during *in utero* growth restriction. The pipeline can be re-iterated to further generate hypotheses (e.g., mechanisms of reversal of impairment), adjust the experimental model (e.g., treatment dosage) and test outcomes. Adaptations of BioRender.com (2021) retrieved from https://app.biorender.com/biorender-templates were made in 1 (adapted from “Mouse Supine”, “Generic cell shape 5, “Placenta” and “Mouse embryo”), 2 (adapted from “Ultrasound transducer (convex)”, “Placenta” and “Blood vessel (3D)”, 4a (adapted from “Capillary bed”) and 5 (“Mouse MRI machine”). Idealized capillaries in 4a were obtained from an open-source dataset^33^ ([Creative Commons Attribution Non Commercial Share Alike 4.0 International](https://creativecommons.org/licenses/by-nc-sa/4.0/legalcode)) and reproduced with permission^31^ (Copyright © 2020, Springer Nature Limited) and fabricated vascular networks in 4b were obtained from an open-source dataset^34^ ([Creative Commons Attribution Non Commercial Share Alike 4.0 International](https://creativecommons.org/licenses/by-nc-sa/4.0/legalcode)) and reproduced with permission^32^ (Copyright © 2019, The American Association for the Advancement of Science).

## Results

### Verification of the Model Against the State-of-the-Art

With the end goal of developing and applying a framework to assess placental function, we began by developing and implementing mathematical models of oxygen diffusion in human placenta capillary-villous structures taken from a seminal placenta oxygen diffusion modelling study ^25^ (Figure 2). We prepared three previously published geometries and ran flow simulations (using parameters listed in Supplementary Material, Table S1) on STAR-CCM+ (v15.02.007) and compared results to Pearce et al. ^25^. Prior to analysis, we ensured that simulation solutions were independent of the discretization approach (See Supplementary Material Table S2, Figure S1). Results were in close agreement for all three geometries tested, with similar wall shear stress profiles (Figure 2A-B) and maximum differences in velocity magnitude below 4.8% (Figure 2C). This suggests that for the low Reynolds numbers typically observed in the capillaries, the finite volume approach to the Navier-Stokes equations used in our model yield similar results to the finite element solution to the Stokes flow, used by Pearce et al. ^25^. Advection-diffusion simulations also yielded visually identical distributions of oxygen concentration (Figure 2D-E) and comparable oxygen transfer rates (< 5%) at the wall interface (Figure 2F). We then used the model to determine new insights into the reduction in oxygen gain with increasing flow rate in the terminal capillary-villous structures, and show that the relationship can be described by negative exponential regressions (Figure 2G).

**Figure 2:**
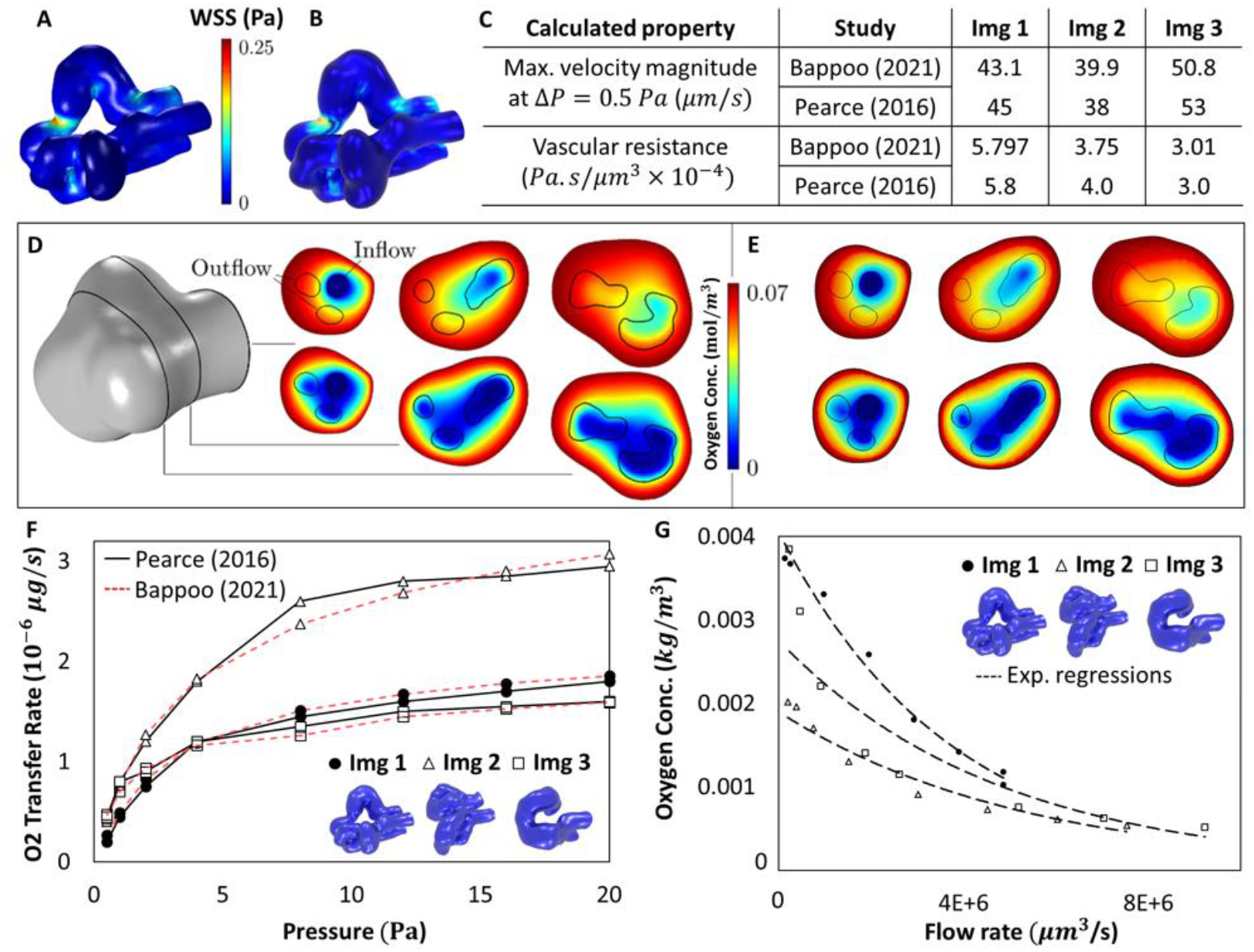
Numerical validation of methods. Flow simulation results derived from CFD methods (full Navier-Stokes equations; STAR-CCM+) used in this study were in close agreement with previous methods (Stokes flow; COMSOL) ^25^. Wall shear stress plots were comparable between ^25^(A) and our methods (B), and other calculated properties (C) were also in close agreement. Oxygen concentration results shown in (D) ^25^ and our results shown in (E) indicate that advection-diffusion results were also in close agreement. Note: We visually approximated the cross-sections presented by Pearce et al.^25^ as these were not indicated. The top row of panels (D) and (E) represent simulations of a pressure drop of 2 Pa while the bottom row was a pressure drop of 20 Pa. The lower pressure drop simulation yielded higher oxygen diffusion into the blood domain. For each human terminal villous specimens (Images 1-3), we show similar behavior of oxygen transfer rate at the tissue-blood interface (F) ^25^, which followed a logarithmic relationship with increasing pressure drop (Image 1: r = 0.9982; Image 2: r = 0.9989; Image 3: r = 0.9987). For each human terminal villous specimen (Images 1-3), we show that higher flow rates reduce the oxygen gained (G), following a negative power exponential relationship (Image 1: r = 0.9939; Image 2: r = 0.9765; Image 3: r = 0.9355). Three-dimensional models of terminal capillary-villi and modelling data (D, F) from Pearce et al.^25^ were adapted under a Creative Commons license ([CC BY 4.0](https://creativecommons.org/licenses/by/4.0/)).

As our goal was to develop a well verified and scalable model for application to more complex geometries, close agreement to the state of the art provided confidence in our approach. Additionally, we demonstrate the ease of scalability as our methods of geometry preparation, computational meshing and solving time using a standard 6GB RAM desktop computer took less than 20 minutes on average, whereas for similar mesh sizes Pearce et al.^25^ required 768GB RAM (computation time was not reported).

### Validation of Oxygen Diffusion Computational Model

Following model verification against state-of-the-art, we aimed to validate the modelling to ensure results represent the ground truth. The gold standard validation approach for computational models are generally *in vivo* experimental measurements, however this is intrinsically difficult to obtain in human placental capillaries due to ethical constraints and the limits of ultrasound, most routinely used in pregnancy. Hence we sourced previously published capillary-inspired geometries and corresponding data of flow-oxygenation experiments (Figure 3) ^32^.

**Figure 3:**
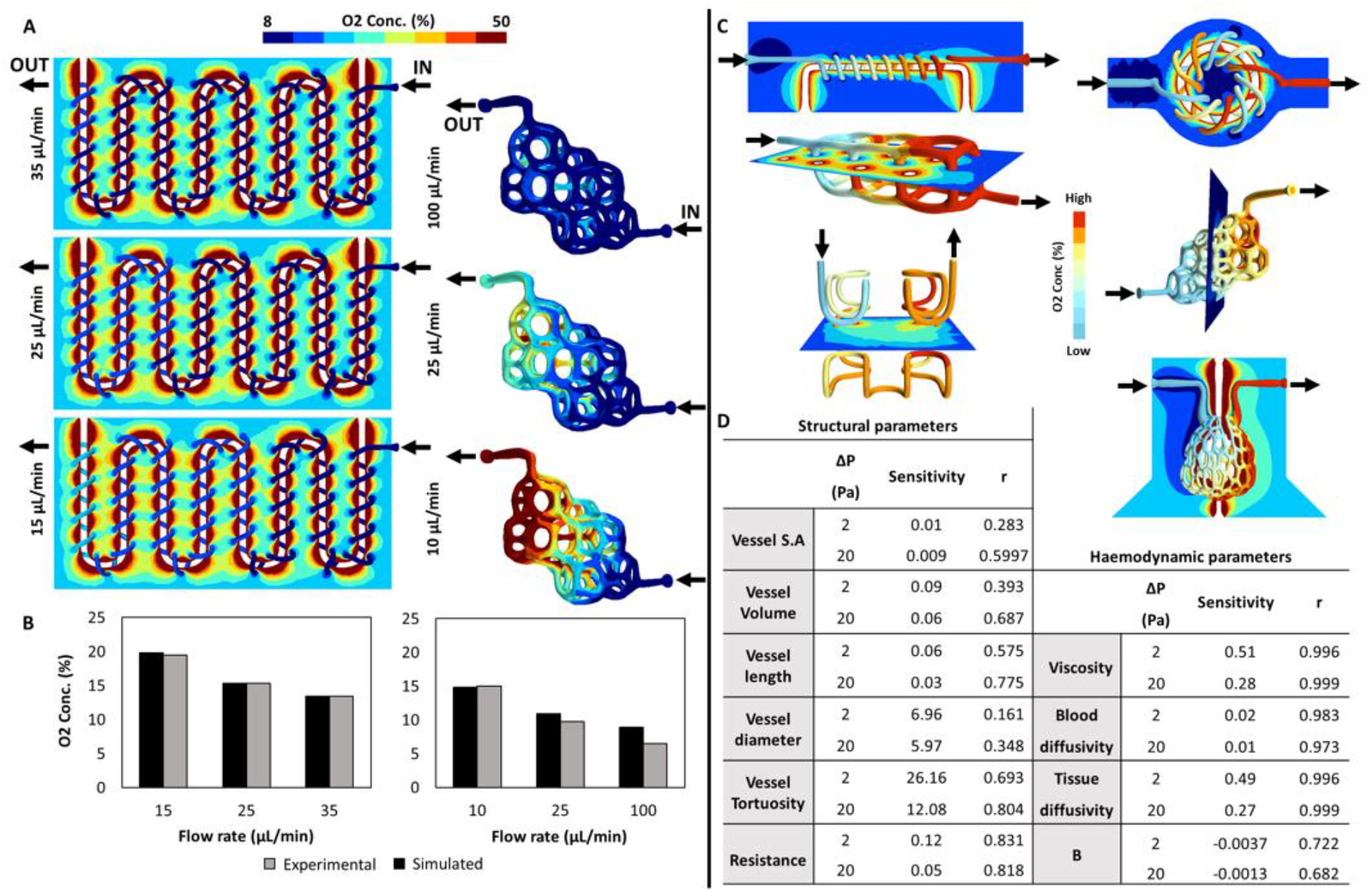
Model development, validation and sensitivity. Calibration and validation model comparison. Contour plots of oxygen concentration are shown for the calibration serpentine-helix geometry (A, *left*) and the validation Weaire-Phelan geometry (A, *right*) simulations at varying inlet flow rates. As the flow rate reduces when progressing through the geometry, higher oxygen gains are observed near the outlets, as shown by higher oxygen concentration. Oxygen concentration gained, as measured at the outlet, is shown for both calibration and validation geometries for both simulation and experimental data for the serpentine-helix geometry (B, left) and Weaire-Phelan geometry (B, right). Model sensitivity was investigated for both structural and haemodynamic parameters was investigated using various geometries (C) and the sensitivity of oxygen gained to both structural and haemodynamic parameters and corresponding Pearson’s correlation coefficients, r, are also shown (D). All geometries (A, C) and experimental data (B) were reproduced with permission (Copyright © 2019, The American Association for the Advancement of Science)^32^ and [Creative Commons Attribution Non Commercial Share Alike 4.0 International](https://creativecommons.org/licenses/by-nc-sa/4.0/legalcode)^34^ .

We firstly used a serpentine-helix geometry (Figure 3A, *right*) for calibration of system unknowns (oxygen diffusivity in hydrogels) using our diffusion methods. See Supplementary Material Figure S2 for further model set up information. We then used results for blind validation of oxygen gain in a Weaire-Phelan geometry (Figure 3A, *left*). The oxygen diffusivity in the gel was calibrated until the oxygen gained reported experimentally in the calibration geometry was matched. We computed the oxygen diffusivity to be 5.81 ×10^−11^ *m*^2^/*s*at the calibration flow rate (35 *μL/min*). We then show that irrespective of flow rates, there was an increase in oxygen concentration as the vasculature progressed from inlet to outlet and that the simulations results were in close agreement with experimental data (Figure 3B, *left*), with errors below 6% recorded (3.4% error at 25 *μL/min*; 5.9% error at 15 *μL/min*). A negative trend was also observed between flow and oxygen gain.

Applying the computed diffusivity to simulations in the Weaire-Phelan geometry (validation geometry) which was manufactured using a similar hydrogel (Note: It was unclear if the same hydrogel was used for both geometries). We showed close agreement to experimental results, as we observed a 1.1% error at 10 *μL/min*, 11.7% error at 25 *μL/min* and 36% error at 100 *μL/min*. (Figure 3B, *right*).

### Sensitivity of the Model to Structural and Haemodynamic Changes

Following model verification and validation, we applied our methods to six previously published topologies of varying complexity ^32^ (Figure 3C) to characterize structural and haemodynamic contributors to oxygen gain. There was an increase in oxygen concentration as the vasculature progressed from inlet to outlet and oxygen gain was pronounced at lower pressure drops. As listed in Figure 3D, structural parameters producing the highest sensitivity were vessel tortuosity (At ΔP = 2 Pa, sensitivity = 26.16; r = 0.69, p = .13. At ΔP = 20 Pa, sensitivity = 12.08; r = 0.80, p= .053), vessel resistance (At ΔP = 2 Pa, sensitivity = 0.12; r = 0.83, p= .82. At ΔP = 20 Pa, sensitivity = 0.05; r = 0.82, p < .05) and vessel diameter, although not significant (At ΔP = 2 Pa, sensitivity = 6.96; r = 0.16, p = .16. At ΔP = 20 Pa, sensitivity = 5.97; r = 0.35, p = .49). Haemodynamic parameters including blood viscosity, blood diffusivity and tissue diffusivity positively correlated with oxygen gain while the advection enhancement factor was negatively correlated. We observed the highest sensitivity for blood viscosity (At ΔP = 2 Pa, sensitivity = 0.51; r = 0.99, p < .01. At ΔP = 20 Pa, sensitivity = 0.28; r = 0.99, p < .01). Detailed results are provided in Supplementary Material (Table S3-5, Figure S3-4).

### Oxygen Diffusion in the Feto-Placental Vasculature

We then applied our oxygen diffusion framework to a well-established rat model of fetal growth restriction, with vascular casting and imaging used to extract pre-capillary arterial feto-placental vascular networks. Limitations of the feto-placental arterial casting method hindered our ability to capture subject-specific placental capillaries in the rat. Human terminal capillaries (Figure 2) were not suitable to couple with the pre-capillary rat placental arterial network due to differences between species ^14^, and bioinspired networks (Figure 3) lacked the complexity of capillaries. Hence, we chose to couple idealized capillaries to downstream vessels of rat feto-placental arteries.

We firstly ensured that our flow simulations produced valid data. To achieve this, we performed computer simulations of an idealized capillary network biofabricated and investigated using particle image velocimetry studies (See Supplementary Material, Figure S5) ^31^. We then simulated oxygen diffusion and developed characteristic curves of oxygen gain by simulating different flow rates in idealized 3D dense and thin capillary networks, for both normoxia (21% oxygen) and hyperoxia (100% oxygen) (Figure 4A-B). In both dense and thin networks, the oxygen gained approached the oxygen source concentration at very low flows and approached zero oxygen gain at very high flows. In both normoxia and hyperoxia scenarios, the dense capillary network was more efficient at gaining oxygen through diffusion.

**Figure 4:**
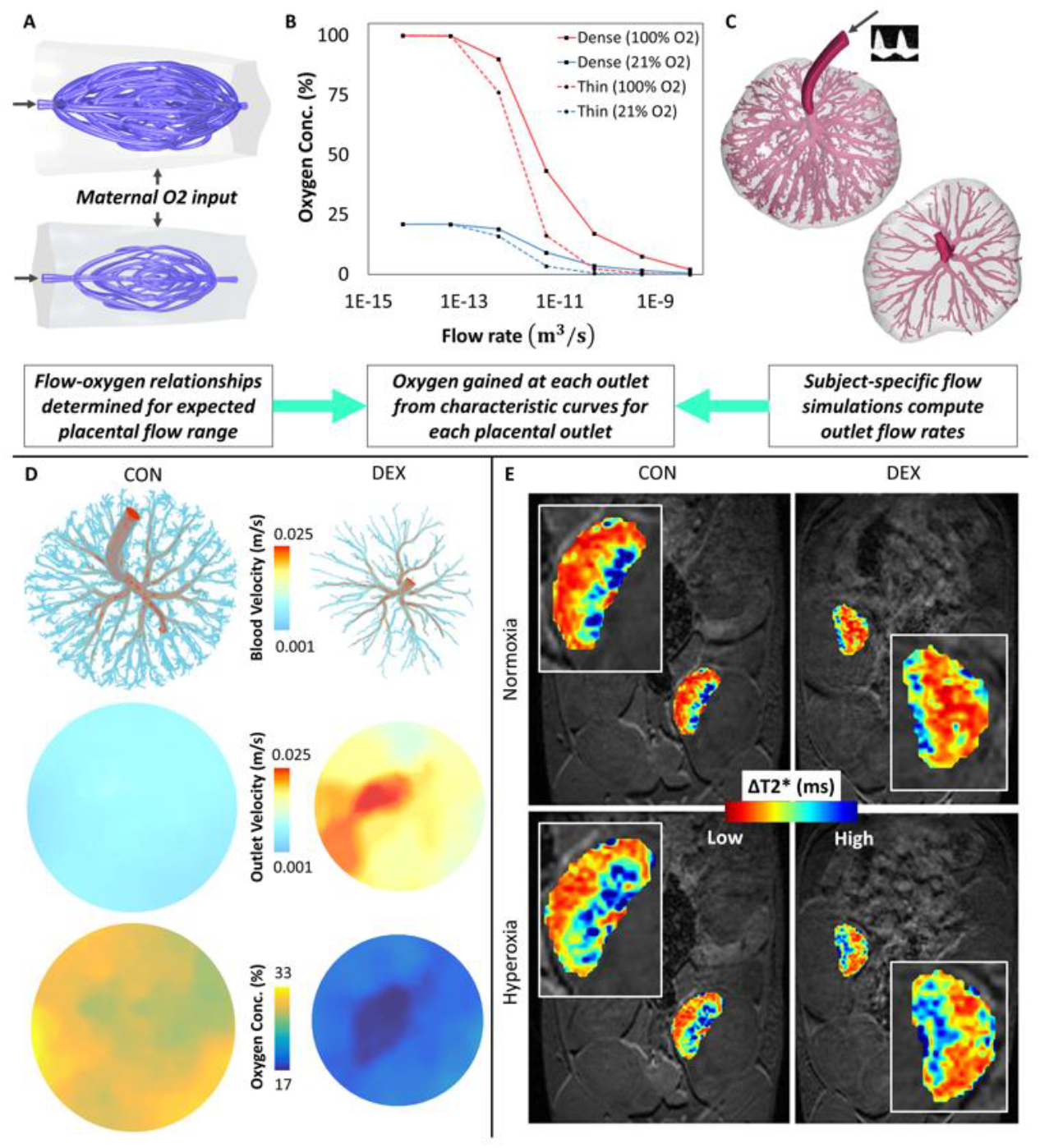
Comparison of flow and oxygenation using computational fluid dynamics (CFD) and MRI. Dense and thin capillary networks (adjusted to the size of feto-placental outlets) were characterized for their flow rate-oxygen behavior (A-B) for maternal oxygen input conditions -normoxia (21% oxygen) and hyperoxia (100% oxygen). Following our previous methods ^13^ high resolution geometries of control (n = 3) and growth-restricted (n = 3) placentas were reconstructed from micro-CT images and converted to a computational mesh for subject-specific CFD simulations (C). Outlet flow rates computed through steady-state simulations interpolate oxygen gain from characteristic curves. We show volumetric renders of blood velocity in control and growth-restricted rat placental vessels, indicating a gradual deceleration of blood from the umbilical artery to the capillary bed (D). This deceleration is more pronounced in control cases as shown by outlet velocities. Oxygen concentrations were computed from characteristic capillary curves (in this case, dense capillaries) and projected onto plane sections for an easily interpretable color map. Derived from MRI (E) are representative 2D slices for normoxia and hyperoxia from the 3D coronal T2* image stacks. Oxygenation was measured through ΔT2* signals (relative change from normoxia to hyperoxia) for control (n = 3) and growth-restricted (n = 3) placentas. Idealized capillaries in (A) were obtained from an open-source dataset^33^ ([Creative Commons Attribution Non Commercial Share Alike 4.0 International](https://creativecommons.org/licenses/by-nc-sa/4.0/legalcode)) and reproduced with permission^31^ (Copyright © 2020, Springer Nature Limited).

Our model of growth restriction resulted in a 21% reduction in fetal weight (p<.0001) and less arborized vascular tree networks (Figure 4C). Growth restricted placentas had 62% fewer vessels (p = .15) and differences in vessel numbers were most pronounced downstream.

Figure 4D shows the example results for the most (control) and least arborized (growth-restricted) placenta. Our simulations revealed a gradual deceleration of blood from the umbilical artery to downstream vessels near the capillary bed in both groups (Figure 4D, first row). However, blood velocity measured at the outlets of the control group was 47% lower (p < .001) than the growth-restricted group. Outlet velocity distributions (Figure 4D, second row) were then projected onto place sections to show key differences in magnitude and heterogeneity. For the case shown in Figure 4D, the growth-restricted case is subject to a localized high outlet velocity region, appearing to be aligned with insertion direction of the umbilical artery.

We then inferred oxygen concentrations from outlet flow rates using the characteristic curves (Figure 4B) and projected these data onto representative plane sections for control and growth-restricted cases (Figure 4D, third row). Regions of high oxygenation corresponded to regions of low outlet velocities in the case shown. Under both hyperoxia and normoxia, oxygenation was approximately 74% higher (p < .05) on average in control cases using the dense capillary model curve and 78% higher (p= .13) for the thin capillary model (Table 1A).

**Table 1:** Relative difference between control and growth-restricted placentas for simulation methods (A) and MRI measurements (B). Simulation results only include the labyrinth zone while results for both the full placenta and labyrinth zone are shown for MRI measurements. Note: Simulation X = idealized dense capillary geometry coupled downstream, Simulation Y = idealized thin capillary geometry coupled downstream.

| A | Relative difference: Control & Dex |  |
| --- | --- | --- |
|  | Simulation X (dense) | Simulation Y (thin) |
| O2 at hyperoxia (100%) | 73.9 | 77.7 |
| O2 at normoxia (21%) | 74.8 | 77.7 |
| B | Relative difference: Control & Dex |  |
|  | Full placenta | Labyrinth zone |
| T2* | 65.3 | 70.1 |

Key to our interpretations of these data was our ability to directly compare observations with equivalent *in utero* imaging data, ΔT2* MRI of the feto-placental domain (Figure 4E), which is an emerging tool to non-invasively quantify placental perfusion and oxygenation ^27^. In control cases, ΔT2* responses computed from the full placenta domain segmentation were approximately 65% higher (p < .0001) or 70% higher (p < .05) for labyrinth zone, indicating a higher level of oxygenation (Table 1B).

We compared the differences (Table 2) between treatment groups computed using our computational model (simulations X and Y) to the MRI data. Comparisons in the labyrinth zone had the least differences (5-10 %), whereas percentages differences ranged from 12-17% in the full domain, which includes the feto-maternal interface.

**Table 2:** Relative difference of control / growth-restricted differences between simulation methods and MRI measurements. Note: Simulation X = idealized dense capillary geometry coupled downstream, Simulation Y = idealized thin capillary geometry coupled downstream.

| A | Relative difference: Simulation X & MRI |  |
| --- | --- | --- |
|  | MRI full domain | MRI labyrinth zone |
| Simulation condition |  |  |
| O2 at hyperoxia (100%) | 12.2 | 5.1 |
| O2 at normoxia (21%) | 13.3 | 6.3 |
| B | Relative difference: Simulation Y & MRI |  |
|  | MRI full domain | MRI labyrinth zone |
| Simulation condition |  |  |
| O2 at hyperoxia (100%) | 16.5 | 9.8 |
| O2 at normoxia (21%) | 16.5 | 9.8 |

## Discussion

Computational modelling of flow and diffusion in vasculature has been an emerging tool in overcoming challenges of traditional qualitative biology and providing novel, often counterintuitive, physical insights into previously unknown mechanisms of both health and disease ^1, 3, 6, 8^. When applied correctly, this tool holds immense diagnostic, prognostic and therapeutic potential in addressing modern era medical conundrums. Although such principles are well established, the complexity of real-world biological systems such as the placenta and lack of validation data represent a methods implementation and data interpretation challenge. Hence, we hypothesized that a simple, scalable and validated *in silico* approach has the potential to overcome limitations of conventional experimental techniques and help generate novel insights on poorly understood structure-function relationships in vascular systems present in exchange organs such as the placenta. Specifically, we hypothesize that the mechanism of functional impairment (exchange) in the growth-restricted group is underlined by abnormally developed vascular structure (hypo-vascularity) which consequently impairs haemodynamic deceleration required for optimal exchange.

Using our innovative framework, we show local and placenta-wide applications of computational modelling complemented by *in utero* 3D imaging and experimental validation data. Through the use of open source mathematical topologies, open-source *ex vivo* specimens of human terminal capillaries, and *in vivo* imaging data from rodent feto-placental vasculature in both health and disease, we have demonstrated that our highly scalable computer model can predict oxygen diffusion.

### Verification and Validation of Our Oxygen Diffusion Computational Model

Our results demonstrated both the ease of implementation and numerical model accuracy across several geometries of varying complexity. Implementation was automated and the overall process per case took less than 20 minutes on average. Our oxygen gain results were in agreement with others, both numerically ^22, 25^ and experimentally ^31, 32^. The calculated oxygen diffusivity in the fabricated gels (*D*_*t*_ = 5.81 ×10^−11^ *m*^2^/*s*) was also consistent with the range of values (*D*_*t*_ = 2.84 ×10^−11^ − 2 ×10^−10^ *m*^2^/*s*) reported in similar hydrogels ^35, 36^.

We acknowledge the larger error observed at the high flow rate (100 *μL/min*) of the validation case which could be a result of changes in the viscosity at higher flows, not accounted for in our computational model. However, low flows are typical of terminal exchange vessels and are therefore more physiologically relevant than higher flows. Alternatively, the properties (diffusivity of oxygen) in the gel used to fabricate the validation case could be different to the calibration case as a result of manufacturing variabilities, hence resulting in a higher error. Although such fabricated gels were used as a validation exercise in this study, our framework has the additional potential to tune and optimize biomaterials and emerging biofabrication methods ^8^ for functional performance.

### Sensitivity of Oxygen Diffusion to Structure and Haemodynamics Changes

Vessel tortuosity was identified as a sensitive structural parameter through our analysis. Tortuosity frequently presents in vascular disease and downstream effects are unknown, however this feature is often ignored in simplified 0D or 1D modelling approaches. For instance, arterial tortuosity has previously been reported as a compensatory remodeling mechanism to increase vessel length and surface area and hence maintain oxygenation in retinopathy ^37^, a theory supported by our results and previous work ^38^. Although seemingly beneficial, we hypothesize that tortuosity is valuable when temporary whereas long term changes are detrimental as they promote blockage, leaky vessels and create an undesirable environment for cells ^39^. Our methods allow for complex geometries with varying structural features to be investigated and downstream physical functionality to be investigated, unlike simplified 0D or 1D approaches.

Interestingly, we also showed that blood viscosity was another highly influential haemodynamic predictor of oxygen diffusion, with effects more pronounced at lower pressure drops (slower flows). It has previously been shown that viscosity increases in microvasculature as a result of the Fåhræus–Lindquist effect ^40^ and this has also been demonstrated in the rat placenta ^13, 18^. We speculate that, macroscopically, this effect (resulting in increased viscosity) aids in local flow deceleration which increases diffusion time, and microscopically, the reduction of the cell-free layer thickness characterized by this effect serves both as a mechanism of flow deceleration and as an adaptation to minimize the distance between red blood cells and tissue to optimize gas exchange. Complex modelling (Discrete Element Method or Lagrangian particles) of deformable blood cells and plasma within the flow domain and incorporation of oxygen-hemoglobin binding mechanisms could test this hypothesis ^41, 42^.

Our methods provide a means to exhaustively actualize physiological theories and define both structural and haemodynamic changes which result in increased oxygen gain. Through our computational framework, features can be manipulated with ease rather than traditional pre-clinical drug or surgical experimentation, opening opportunities to devise, optimize and maximize value of pre-clinical *in vitro* and animal models.

### Oxygen Diffusion in the Feto-Placental Vasculature is Dependent on Structure and Flow Deceleration

Growth-restricted rodent feto-placental networks resulted in a markedly less arborized vascular network. We hypothesized that a larger number of vessels is necessary for optimal flow deceleration and ultimately, oxygen diffusion. Our data confirmed that this behavior, finding control feto-placental vascular networks were more efficient at decelerating blood compared to growth-restricted placentas. Murray’s Law has previously been used to explain that the development of expansive hierarchical bifurcating tree structures is necessary to ensure that blood flows away from the heart with gradual pressure loss ^43^, which results in significant flow deceleration and increased time for exchange to efficiently occur at the capillary level.

The placenta develops extensive vasculature to achieve its function of exchanging gases, nutrients and waste, hence it becomes evident that any structural impairment would impair exchange, yet not clearly demonstrated. Here we present the novel use of idealized capillary models coupled with the feto-placental vessels to simulate downstream flows and consequential diffusion behavior. We characterized flow rate-oxygenation behavior in idealized capillaries and showed the oxygen levels approached the source concentration at very low flows and conversely, diffusion became negligible at very high flows. Indeed, when coupled with downstream velocities of feto-placental structures, the higher outlet blood velocities modelled in growth-restricted placentas led to impaired oxygen exchange. Key to our interpretation and confidence in these simulation results were our MRI validation cohort which also showed impaired oxygenation in growth-restriction (70% lower). We demonstrated the simulation assumption of idealized dense capillaries led to the least difference to MRI measurements in the labyrinth zone (< 7%) compared to thin capillaries (< 10%). We also show an example of a full placenta with no idealized capillaries coupled in Supplementary Material (Table S6, Figure S6) (< 9% difference compared to MRI data), however this approach assumes non-physiological diffusion to occur in larger pre-capillary vessels. These data represent a major advancement to this area of research and to our knowledge, has not been reported elsewhere. Not only has the *in vivo* image data validated our computational approach, it also demonstrates that when used in tandem, these emerging interdisciplinary tools can overcome current challenges of traditional biology, and may enable future novel advancements.

This is the first study to demonstrate the feasibility of computational fluid dynamics (flow and diffusion) as a reliable and well-validated tool in feto-placental vasculature networks. The pipeline overcomes current challenges in implementing complex computational methods in the placenta and the difficulty of model validation ^30^ to demonstrate localised effects of impaired vascular structure and consequently function (oxygen exchange). We hypothesized that the mechanism of functional impairment (exchange) in the growth-restricted group is underlined by abnormally developed vascular structure (hypo-vascularity) which in turn impairs blood flow deceleration, and hence reduces the transit time for optimal exchange. Expanding our readily implementable methods to a statistically powered experimental cohort would confirm this hypothesis.

## Limitations

Although the idealized capillary assumption produced replicable and validated results, we acknowledge that feto-placental capillaries *in utero* are unlikely to perfectly resemble our bioinspired models. Additionally, the closest agreement between computational simulations and MRI measurements was shown when the dense capillary model was coupled to both control and growth restricted placentas, but physiologically, dense capillaries should only be assumed in control cases (and thin capillaries in growth restriction) as it has previously been shown that glucocorticoid exposure reduces downstream fetal capillaries ^15, 17^. Hence, advancements in casting, imaging methods ^44-46^ and segmentation of capillary vessels will lead to better *in silico* models.

A limitation to our current experimental validation approach is the spatial resolution of MRI (0.5 mm x 0.5 mm x 0.5 mm), which upon improvement, would help distinguish between intravascular and tissue oxygenation and improve interpretation of results to a similar detail as the computational model. Here, we compare percentage differences between control and growth-restricted groups computed using either our computer simulations (absolute total concentration at either hyperoxia or normoxia) or MRI (ΔT2* response from normoxia to hyperoxia). An improved and more direct validation of our simulations would involve comparing absolute oxygen values measured from both simulation and MRI methods.

However, absolute measurements are not currently feasible using the MRI method which relies on a T2* response measurement. Conversely, simulating the T2* response would involve a range of assumptions including maternal responses to the oxygen challenge or placental and fetal metabolism, which would increase simulation complexity and introduce unnecessary unknowns. Finally, although differences between computational and MRI results were minimal, further validation in a larger cohort is needed to establish our proposed mechanism of impairment.

## Future Perspectives

We envision our framework to further dissect complex biological phenomena while creating opportunities for blue-sky research and development of novel technologies in image-and simulation-based diagnostics or prognostics, an area which has recently gained momentum through FDA-approved products^47^. In the first instance, we aim to apply the framework to study human placentas, however there are opportunities to study tissue oxygenation in other systems such as the retina or tumors and to incorporate additional complexity such as tissue metabolism or active transport to study advanced drug delivery^6^.

## Methods

### Geometries

All geometries except rat feto-placental arterial cases presented, were obtained from open-source publication data, particularly bioinspired intravascular topologies ^31, 32^ and feto-placental capillary-villi images ^25^.

#### Human placenta capillary-villous structures

Specimens of terminal human placenta capillary-villous structures imaged with confocal laser scanning microscopy (Figure 2) were obtained through an open-source publication ^25^ and used for numerical validation of our methods.

#### Synthetic vascular networks

To calibrate the computational model, we used an entangled serpentine-helix network (Figure 3A, *left*), available as triangulated iso-surfaces in stereo lithography (stl) format from Grigoryan et al.^32^ who studied blood oxygenation through a 3D bioprinted aqueous hydrogel (20 wt%, 6-kDa PEGDA) and reported measured oxygen concentrations. We then used a Weaire-Phelan vascular network (Figure 3A, *right*) ^32^ to validate the calibrated model. Other topologies (Figure 3C) from the same open-source publication were used to test the sensitivity of the model to varying structures. We also sourced idealized dense and thin capillary networks (Figure 4A)^31^ to couple with near-capillary vessels in placental applications, described later.

#### Animal ethics

All animal studies were performed according to animal health and welfare guidelines and were approved by the Animal Ethics Committee of the University of Western Australia (AEC number RA/3/100/1600). Wistar rats (age 6-8 weeks old) were purchased from the Animal Resources Centre (Murdoch, WA, Australia) and housed under a 12 hour light dark cycle. Food and water were available *ad libitum*, except during brief testing periods.

#### Rat feto-placental vascular structures

We collected, imaged and segmented rat feto-placental vasculature using our methods previously described ^13, 18, 20, 48^. Briefly, female Wistar rats were time-mated and administered with either vehicle (control group, n=3) or 0.5 µg/ml dexamethasone acetate as a model of growth restriction in fetuses (growth-restricted group, n=3). Feto-placental arterial vasculature from both groups were then cast towards the end of gestation (Embryonic Day 21, E21) and solidified casts were scanned using micro-CT (Xradia 520 Versa X-ray Microscope, ZEISS, Oberkochen, Germany) (Figure 4C).

## Computational Model

We modelled steady fluid flow in STAR-CCM+ using a three-dimensional (3D) finite-volume discretization of the incompressible Navier-Stokes and continuity equations using established methods ^13, 18^. Furthermore, a passive scalar formulation was used to solve for steady-state solute (oxygen) concentration throughout the fluid (i.e., blood, water) and solid (i.e. tissue, gel media) regions, using the advection-diffusion equation (Eq. 1).

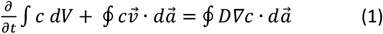

where, *c* is the concentration, 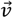 is the fluid velocity and *D* is the diffusivity.

For steady-state conditions, the transport of the solute concentration c in the fluid and solid media may be expressed by Eq. 2 and Eq. 3, respectively, where no advection occurs in the solid media.

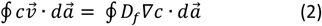

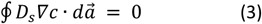

where, *D*_*f*_ is the diffusivity of the solute in the fluid and *D*_*s*_ is the diffusivity of the solute in the solid.

To account for the binding of oxygen to hemoglobin when modelling the transport of oxygen in blood, an additional oxygen advection enhancement parameter (*B*) was incorporated through coupling with the velocity field in the fluid domain ^22, 23, 25^ (Eq. 4).

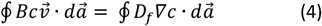

The numerical solution of the advection-diffusion equation was performed after the fluid velocity field reached steady-state: the root-mean-square (RMS) value of the absolute momentum and continuity residuals were below 10^−13^.

Simulated flow was validated experimentally using open-source publication data ^31^ which used particle image velocimetry (PIV) for model comparison. Results were in close agreement and further information can be found in Supplementary Material (Figure S3).

### Discretization Approach

The computational model consisted of two domains in all geometries simulated; a blood (fluid) domain embedded within a gel or tissue (solid) domain. We discretized both of those regions with trimmer cells and prism-layer cells, however prism-layers in the solid domain were only assigned at the near-wall boundary adjacent to the blood domain. Numerical accuracy in both domains was confirmed using a Grid Convergence Index (GCI) test ^49, 50^, which ensures that computed solutions are independent of the discretization approach. Further information is provided in Supplementary Material (Table S2; Figure S1).

### Boundary conditions

All geometries had one inlet and one outlet boundary, except for rat feto-placental arterial geometries that had one inlet and multiple outlets (ranging from 199-1406 outlets).

Generally, simulations were set up as illustrated in Figure 1, with input boundary conditions assigned at the inlet and at oxygen sources. At the internal interface, the solute concentration and diffusive solute fluxes were equal. For calibration (serpentine-helix), we assigned parabolic velocity profiles matching experimental flow rates at the inlet, 35, 25 *and* 15 *μL/min*, while flow rates were 10, 25 *and* 100 *μL/min* for the validation case (Weaire-Phelan). We measured the oxygen concentration at the outlet as volumetric concentrations. For sensitivity analyses and numerical model comparisons, a similar approach using pressure drops was used to simulate flow, rather than prescribed flow rates.

### Model assumptions

For all geometries, the system was assumed to be closed, hence conservation of mass was imposed. We assumed rigid walls with a no-slip condition defined.

We assumed constant network hematocrit and adjusted the viscosity to account for body temperature (37°C) and low shear rates in the simulated flows. The internal oxygen source (gas domain interface in Figure S2) was assumed to be a 100% oxygen source over its entire domain as experimental oxygen conditions reported by Grigoryan et al.^32^ were not quantitatively specified. We assumed that experiments were performed at standard room conditions, hence at the gel-air interface, we assumed a fixed concentration of 21% oxygen ^51^. In these cases where fetal blood was not used, the oxygen advection enhancement parameter was assumed to be 1, otherwise, the reader may refer to Pearce et al.^25^ for detailed subject-specific calculations in cases of fetal blood.

For our comparison to a previous numerical model, parameters in Table 3 were adjusted to values used by Pearce et al.^25^, available in Supplementary Material (Table S1). Rat feto-placental arterial simulations followed assumptions previously reported, which account for differences in species and scale ^13^. However, blood and tissue diffusivity were assumed as values in Table 3.

**Table 3:** Literature based parameters used in the mathematical model.

| Parameter | Symbol | Value | Ref. |
| --- | --- | --- | --- |
| Dynamic blood viscosity at 37°C | $\mu$ | 0.007 Pa.s | 52 |
| Diffusivity of oxygen in plasma | $D_p$ | $1.7 \times 10^{-9} \text{ m}^2/\text{s}$ | 53 |
| Diffusivity of oxygen in hydrogel | $D_t$ | Calculated | N/A |

### Model Calibration and Validation

Using the serpentine-helix geometry (Figure 2A, left) and available experimental data of flow and oxygenation, the computational model was calibrated to match experimental conditions. Firstly, the experimental values of inlet and outlet *S*_*0*2_ (%) and *P*_*0*2_ (*mmHg*) were converted to a volumetric concentration using Eq. 5 and expressed as a percentage. The inlet oxygen concentration was derived as 8.22 % (*S*_*0*2_ = 41.4%; *P*_*o*2_ = 33.5%) while the target outlet oxygen concentration was 13.41 % (*S*_*0*2_ = 67.3%; *P*_*0*2_ = 70.4%).

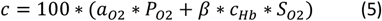

where *β* is the oxygen capacity of blood, assumed to be 1.4 *mL · g*^−1 54^, *a*_*o*2_ is the proportionality constant derived from Henry’s Law for oxygen in blood, assumed to be 0.0031 *mL · mmHg*^−1^*dL*^−1 55^ and *c*_*Hb*_ is the mass concentration of haemoglobin in blood, assumed to be 14 *g · dL*^−1 54^.

The calibration simulation was then run at an inlet flow rate of 35 *μL/min* and the oxygen diffusivity of the gel (or tissue) *D*_*t*_, an unknown to the system, was gradually iterated until the computational model produced results matching the experimental data, the oxygen concentration at the model outlet, expressed as volumetric concentrations. The computed *D*_*t*_ value was verified against similar hydrogels reported in the literature. Once calibrated, the computed *D*_*t*_ value was applied to the validation Weaire-Phelan model and a blind comparison to experimental values was made. Simulations were also run at other flow rates in both geometries to further validate flow-diffusion relationships shown experimentally ^32^.

## Sensitivity analysis

Following calibration and validation of methods presented, we performed a sensitivity analysis (Figure 3C-D) on the computational model to reveal the influence of parameters on oxygen gain, by incrementally varying original values used (Table 3). Parameters included blood viscosity, blood and tissue diffusivity and the advection enhancement term and were normalized to values in Table 3. The computed outlet oxygen concentration were also normalized to the computed concentration when values in Table 3 were used. Sensitivity of outlet oxygen concentration to parameters was computed through least squares linear regressions.

We also investigated the impact of structure on oxygen diffusion. Bioinspired vascular geometries ^32^ (Figure 3C) were used and pressure drops of 2 Pa and 20 Pa were simulated across fluid domains. Structural parameters such as vessel and gel surface area and volume, vessel centerline length, vessel diameter and vessel tortuosity were extracted on Mimics (Materialise, Belgium). Vessel resistance was calculated as the ratio of pressure drop to volumetric flow rate across the vessel network. As before, sensitivity of the oxygen gain to structural parameters were calculated.

### Numerical Method Comparison

Pearce et al.^25^ implemented Stokes flow as well as advection-diffusion equations on COMSOL Multiphysics (finite-element), instead of our approach using the Navier-Stokes equations in STAR-CCM+ (finite-volume). We used three human terminal placental capillary to compare numerical methods under similar conditions as reported^25^ (Table S1). From our simulation, we compared shear stress contours (Figure 2A-B) and maximum velocities (Figure 2C). We also compared contour plots of oxygen distribution in 3 planes approximately similar to those presented in Pearce et al.^25^ (Figure 2D-E) and compared the total oxygen transfer rate, calculated by integrating the advective flux leaving the capillary domains at the outlets (Figure 2F). Finally, we investigated the relationship between total oxygen gained and the inlet flow rate to demonstrate diffusion behavior in *ex vivo* placental specimens (Figure 2G).

### Feto-placental Blood Flow Simulations

For placenta simulations, we reconstructed 3D high resolution geometries of control and growth-restricted rat feto-placental arteries collected through a polymer casting method and micro-CT methods described previously. High frequency Doppler ultrasound was used to measure the umbilical artery velocity using established methods ^16^ for each group for use as the model input (waveform integrated over the inlet to simulate mean flow conditions) and asymmetrical structured fractal trees at outlets were used to model the resistance of unresolved peripheral vessels ^13^. Simulations were run using well established methods and outlet flow rates were recorded.

Gas diffusion was assumed to occur downstream of the truncated feto-placental vessels, however rat feto-placental capillaries could not be routinely resolved using our casting and imaging methods ^56^. Hence we aimed to utilize computed outlet flow rates (from placental simulations, Figure 4C) as inputs to representative open-source capillary geometries ^31^ (Figure 4A).

Briefly, we first adjusted the size of the open-source geometries by scaling the vessel inlet area of the geometry to match the average feto-placental outlet area. We ran flow simulations in a dense and a thin capillary network with capillary inlet flow rates in the range of flow rates captured from outlets of full feto-placental network simulations and created characteristic flow-oxygen gain curves (Figure 4B). We also compared results for two maternal oxygen input conditions, at normoxia (21% oxygen) and hyperoxia (100% oxygen) for comparison with the maternal oxygen challenge T2* MRI experiments described later. Once characterized, the flow-oxygen curves were splined on Matlab (R2017b) and outlet flow rates recorded from full placental simulations described above were used to interpolate a corresponding oxygen value from the characteristic curve.

In summary, two simulation configurations were investigated. In each configuration, the oxygen gained was measured for control (n = 3) and growth-restricted (n = 3) cases and the relative difference between treatment groups was calculated (Figure 4D). This relative treatment group difference from the various simulation scenario results was compared to the MRI results for validation.

1. **Simulation Condition X:** Characteristic flow rate-oxygen curve of dense capillary network (in which diffusion occurs) used to calculate oxygen gain for each outlet of both control and growth-restricted placentas and for both normoxia (21% oxygen) and hyperoxia (100% oxygen).
2. **Simulation Condition Y:** Characteristic flow rate-oxygen curve of thin capillary network (in which diffusion occurs) used to calculate oxygen gain for each outlet of both control and growth-restricted placentas and for both normoxia (21% oxygen) and hyperoxia (100% oxygen).

### Feto-placental MRI

Similar to the feto-placental vascular casting protocol described previously, pregnant rats from control and growth-restricted groups were imaged on the last day of gestation (E21). All MRI scans were performed under general anesthesia and the animal’s temperature and respiration rate were monitored using a PC-SAM Small Animal Monitor. The temperature of the cradle and the anesthetic gas mixture were adjustable during the MRI scan if required to maintain stable physiology.

A 9.4 Tesla (T) MRI scanner (Bruker BioSpec 94/30 US/R magnet) with BGA-12SHP gradients, AVANCE III HD console, ParaVision 6.0.1 software, and 86 mm ID quadrature H-1 volume coil was used to capture 3D *in utero* measurements of feto-placental oxygenation.

As baseline conditions of T2* are variable due to extraneous factors (e.g., breathing motion, maternal or fetal movements), the measured absolute T2* signal from the placenta is not directly representative of an absolute value of placental oxygenation.

Instead, a maternal oxygen challenge was used as an alternative to capture relative changes irrespective of baseline conditions and extraneous factors. Differences in T2* (ΔT2*) values across the placental membrane during an oxygen-challenge (from normoxia or medical air (21% oxygen) to hyperoxia or 100% medical oxygen) were therefore used to capture the blood oxygen concentration gradient, and thus oxygen transfer across the placenta ^57, 58^ Using the T2-weighted data, multiple 2D regions-of-interest (ROIs) were drawn manually in three planes (coronal, axial, and sagittal). T2* values for the oxygen-challenge paradigm were calculated using custom-written Matlab® scripts based on the SQEXP algorithm ^59^. The ROIs in three planes were drawn for whole placentas and the labyrinth zone; the site of maternal-fetal exchange of the rat placentas. The ROIs were overlaid onto the T2*-weighted gradient-echo image at echo time 5.5 ms (Figure 4E). For further detail on the MRI experimental protocols, refer to Supplementary Material provided (Table S7).

ΔT2* was measured for control and growth-restricted cases and the relative difference between treatment groups was calculated. This relative treatment group difference from the MRI was compared to the various simulation scenario results.

### Statistics

Regressions used were calculated on Excel using in-built least squares regression methods and Pearson’s correlation coefficient was also calculated for normally distributed data (assessed through a Kolmogorov-Smirnov Test of Normality). Linear regressions were used for model sensitivity analyses, while exponential regressions were applied to (i) oxygen concentration-flow rate relationships in terminal human placental capillaries.

Characteristic flow rate-oxygen curves of open-source dense and thin capillary networks were splined using the Matlab 2019b curve fitting tool and flow rates computed in full network simulations of control and growth-restricted placentas were used to interpolate an oxygen value for both normoxia and hyperoxia conditions. Computed outlet velocities and oxygen concentrations derived from either dense or thin capillary curves, were projected onto a plane section for the visualization of downstream blood flow behavior.

The relative difference between total oxygen gained by the growth-restricted placentas with respect to control cases was calculated at both normoxia and hyperoxia. The T2* equivalent response (from normoxia to hyperoxia) was not simulated due to complex underlying modelling limitations discussed later, therefore simulation results at normoxia and hyperoxia were directly compared to T2* MRI (measure of oxygenation) of the feto-placental networks.

It is also important to know that although we report p-values for our analyses, low sample sizes (n = 3 per group) were used. Normally distributed variables were analysed using Student’s t-test while non-normal distributed variables were analysed with Mann-Whitney U test. Correlations for linearly associated variables were analysed using Pearson’s Correlation Coefficient. Differences were accepted as statistically significant when p < .05.

## Acknowledgements

We thank the large number of open-source and related projects that facilitated this work, including the three-dimensional images of terminal human capillary-villi made available by Pearce et al. ^25^ (adapted under a Creative Commons licence ([CC BY 4.0](https://creativecommons.org/licenses/by/4.0/))) and three-dimensional images of multivascular networks and accompanying experimental data made available by Grigoryan et al.^32^ (Copyright © 2019, The American Association for the Advancement of Science) and Miller et al.^34^ ([Creative Commons Attribution Non Commercial Share Alike 4.0 International](https://creativecommons.org/licenses/by-nc-sa/4.0/legalcode)) as well as Kinstlinger et al.^31^ (Copyright © 2020, Springer Nature Limited) and Miller et al.^33^ ([Creative Commons Attribution Non Commercial Share Alike 4.0 International](https://creativecommons.org/licenses/by-nc-sa/4.0/legalcode)). We would like to acknowledge funding from the National Health and Medical Research Council (Grants APP1083572) and the Department of Health, Government of Western Australia. We would also like to acknowledge the facilities, and the scientific and technical assistance of both the National Imaging Facility at the Centre for Microscopy, Characterization and Analysis at The University of Western Australia (a facility funded by the University, State and Commonwealth Governments). This work was also supported by resources provided by the Pawsey Supercomputing Centre with funding from the Australian Government and the Government of Western Australia.

## Author Contributions

NB was involved in planning the study, formulating the modelling methods, analyzing the structural and haemodynamic data and was the main writer of the manuscript. YT was involved in the animal work, vascular casting, and MRI experiments. LJK was involved in developing and optimizing the methods for computational modelling. KWF contributed to the development of MRI protocols. HC contributed to numerical methods verifications. CW and BJD contributed to the study conception, analyzing the structural and haemodynamic data, and discussion and interpretation of the data. CW and BJD supervised the research and take overall responsibility for the work. All authors contributed to the preparation and approval of the final submitted manuscript.

## References

1. Lins M, Vandevenne J, Thillai M, Lavon BR, Lanclus M, Bonte S, Godon R, Kendall I, De Backer J and De Backer W. Assessment of Small Pulmonary Blood Vessels in COVID-19 Patients Using HRCT. Acad Radiol. 2020;27:1449–1455.

2. Donovan GM, Langton D and Noble PB. Phenotype-and patient-specific modelling in asthma: Bronchial thermoplasty and uncertainty quantification. J Theor Biol. 2020;501:110337.

3. Clark AR, Lee TC and James JL. Computational modeling of the interactions between the maternal and fetal circulations in human pregnancy. Wires Syst Biol Med. 2021;13.

4. Jensen OE and Chernyavsky IL. Blood Flow and Transport in the Human Placenta. Annu Rev Fluid Mech. 2019;51:25–47.

5. Mayo RP. Advances in Human Placental Biomechanics. Computational and Structural Biotechnology Journal. 2018;16:298–306.

6. d’Esposito A, Sweeney PW, Ali M, Saleh M, Ramasawmy R, Roberts TA, Agliardi G, Desjardins A, Lythgoe MF, Pedley RB, Shipley R and Walker-Samuel S. Computational fluid dynamics with imaging of cleared tissue and of in vivo perfusion predicts drug uptake and treatment responses in tumours. Nat Biomed Eng. 2018;2:773–787.

7. Altrock PM, Liu LL and Michor F. The mathematics of cancer: integrating quantitative models. Nat Rev Cancer. 2015;15:730–45.

8. Lovett M, Lee K, Edwards A and Kaplan DL. Vascularization strategies for tissue engineering. Tissue Eng Part B Rev. 2009;15:353–70.

9. Liu D, Wood NB, Witt N, Hughes AD, Thom SA and Xu XY. Computational analysis of oxygen transport in the retinal arterial network. Curr Eye Res. 2009;34:945–56.

10. Menshykau D, Michos O, Lang C, Conrad L, McMahon AP and Iber D. Image-based modeling of kidney branching morphogenesis reveals GDNF-RET based Turing-type mechanism and pattern-modulating WNT11 feedback. Nat Commun. 2019;10:239.

11. Buchwald P. FEM-based oxygen consumption and cell viability models for avascular pancreatic islets. Theor Biol Med Model. 2009;6:5.

12. Burton GJ and Jauniaux E. Sonographic, stereological and Doppler flow velocimetric assessments of placental maturity. Br J Obstet Gynaecol. 1995;102.

13. Bappoo N, Kelsey LJ, Tongpob Y, Wyrwoll C and Doyle BJ. Investigating the Upstream and Downstream Hemodynamic Boundary Conditions of Healthy and Growth-Restricted Rat Feto-Placental Arterial Networks. Ann Biomed Eng. 2021.

14. Georgiades P, Ferguson-Smith AC and Burton GJ. Comparative developmental anatomy of the murine and human definitive placentae. Placenta. 2002;23.

15. Hewitt DP, Mark PJ and Waddell BJ. Glucocorticoids prevent the normal increase in placental vascular endothelial growth factor expression and placental vascularity during late pregnancy in the rat. Endocrinology. 2006;147:5568–74.

16. Wyrwoll CS, Noble J, Thomson A, Tesic D, Miller MR, Rog-Zielinska EA, Moran CM, Seckl JR, Chapman KE and Holmes MC. Pravastatin ameliorates placental vascular defects, fetal growth, and cardiac function in a model of glucocorticoid excess. Proc Natl Acad Sci U S A. 2016;113:6265–70.

17. Wyrwoll CS, Seckl JR and Holmes MC. Altered placental function of 11beta-hydroxysteroid dehydrogenase 2 knockout mice. Endocrinology. 2009;150:1287–93.

18. Bappoo N, Kelsey LJ, Parker L, Crough T, Moran CM, Thomson A, Holmes MC, Wyrwoll CS and Doyle BJ. Viscosity and haemodynamics in a late gestation rat feto-placental arterial network. Biomech Model Mechanobiol. 2017;16:1361–1372.

19. Rennie MY, Detmar J, Whiteley KJ, Yang J, Jurisicova A, Adamson SL and Sled JG. Vessel tortuousity and reduced vascularization in the fetoplacental arterial tree after maternal exposure to polycyclic aromatic hydrocarbons. Am J Physiol Heart Circ Physiol. 2011;300:H675–84.

20. Tongpob Y, Xia S, Wyrwoll C and Mehnert A. Quantitative characterization of rodent feto-placental vasculature morphology in micro-computed tomography images. Computer Methods and Programs in Biomedicine. 2019;179:104984.

21. Yang J, Yu LX, Rennie MY, Sled JG and Henkelman RM. Comparative structural and hemodynamic analysis of vascular trees. Am J Physiol Heart Circ Physiol. 2010;298:H1249–59.

22. Erlich A, Pearce P, Mayo RP, Jensen OE and Chernyavsky IL. Physical and geometric determinants of transport in fetoplacental microvascular networks. Sci Adv. 2019;5:eaav6326.

23. Erlich A, Nye GA, Brownbill P, Jensen OE and Chernyavsky IL. Quantifying the impact of tissue metabolism on solute transport in feto-placental microvascular networks. Interface Focus. 2019;9:20190021.

24. Nye GA, Ingram E, Johnstone ED, Jensen OE, Schneider H, Lewis RM, Chernyavsky IL and Brownbill P. Human placental oxygenation in late gestation: experimental and theoretical approaches. J Physiol. 2018;596:5523–5534.

25. Pearce P, Brownbill P, Janacek J, Jirkovska M, Kubinova L, Chernyavsky IL and Jensen OE. Image-Based Modeling of Blood Flow and Oxygen Transfer in Feto-Placental Capillaries. PLoS One. 2016;11:e0165369.

26. Clark AR, Lin M, Tawhai M, Saghian R and James JL. Multiscale modelling of the feto-placental vasculature. Interface Focus. 2015;5:20140078.

27. Aughwane R, Ingram E, Johnstone ED, Salomon LJ, David AL and Melbourne A. Placental MRI and its application to fetal intervention. Prenatal Diag. 2019.

28. Avni R, Neeman M and Garbow JR. Functional MRI of the placenta--From rodents to humans. Placenta. 2015;36:615–22.

29. Ingram E, Morris D, Naish J, Myers J and Johnstone E. MR Imaging Measurements of Altered Placental Oxygenation in Pregnancies Complicated by Fetal Growth Restriction. Radiology. 2017;285:953–960.

30. Byrne M, Aughwane R, James JL, Hutchinson JC, Arthurs OJ, Sebire NJ, Ourselin S, David AL, Melbourne A and Clark AR. Structure-function relationships in the feto-placental circulation from in silico interpretation of micro-CT vascular structures. J Theor Biol. 2021;517:110630.

31. Kinstlinger IS, Saxton SH, Calderon GA, Ruiz KV, Yalacki DR, Deme PR, Rosenkrantz JE, Louis-Rosenberg JD, Johansson F, Janson KD, Sazer DW, Panchavati SS, Bissig KD, Stevens KR and Miller JS. Generation of model tissues with dendritic vascular networks via sacrificial laser-sintered carbohydrate templates. Nat Biomed Eng. 2020;4:916–932.

32. Grigoryan B, Paulsen SJ, Corbett DC, Sazer DW, Fortin CL, Zaita AJ, Greenfield PT, Calafat NJ, Gounley JP, Ta AH, Johansson F, Randles A, Rosenkrantz JE, Louis-Rosenberg JD, Galie PA, Stevens KR and Miller JS. Multivascular networks and functional intravascular topologies within biocompatible hydrogels. Science. 2019;364:458–464.

33. Miller J. Dataset for: Generation of model tissues with dendritic vascular networks via sacrificial laser-sintered carbohydrate templates. 2020.

34. Miller J. Dataset for: Multivascular networks and functional intravascular topologies within biocompatible hydrogels. 2019.

35. An HZ, Eral HB, Chen L, Chen MB and Doyle PS. Synthesis of colloidal microgels using oxygen-controlled flow lithography. Soft Matter. 2014;10:7595–7605.

36. Debroy D, Oakey J and Li D. Interfacially-mediated oxygen inhibition for precise and continuous poly(ethylene glycol) diacrylate (PEGDA) particle fabrication. J Colloid Interface Sci. 2018;510:334–344.

37. Khansari MM, Garvey SL, Farzad S, Shi Y and Shahidi M. Relationship between retinal vessel tortuosity and oxygenation in sickle cell retinopathy. Int J Retina Vitreous. 2019;5:47.

38. Goldman D and Popel AS. A computational study of the effect of capillary network anastomoses and tortuosity on oxygen transport. J Theor Biol. 2000;206:181–94.

39. Szabo A and Merks RMH. Blood vessel tortuosity selects against evolution of aggressive tumor cells in confined tissue environments: A modeling approach. PLoS Comput Biol. 2017;13:e1005635.

40. Pries AR, Secomb TW, Gebner T, Sperandio MB, Gross JF and Gaehtgens P. Resistence to blood flow in microvessels in vivo. Circ Res. 1994;75.

41. Tsai AG, Johnson PC and Intaglietta M. Oxygen gradients in the microcirculation. Physiol Rev. 2003;83:933–63.

42. Ong PK, Namgung B, Johnson PC and Kim S. Effect of erythrocyte aggregation and flow rate on cell-free layer formation in arterioles. Am J Physiol Heart Circ Physiol. 2010;298:H1870–8.

43. Murray CD. The Physiological Principle of Minimum Work: I. The Vascular System and the Cost of Blood Volume. Proc Natl Acad Sci U S A. 1926;12:207–14.

44. Cai R, Pan C, Ghasemigharagoz A, Todorov MI, Forstera B, Zhao S, Bhatia HS, Parra-Damas A, Mrowka L, Theodorou D, Rempfler M, Xavier ALR, Kress BT, Benakis C, Steinke H, Liebscher S, Bechmann I, Liesz A, Menze B, Kerschensteiner M, Nedergaard M and Erturk A. Panoptic imaging of transparent mice reveals whole-body neuronal projections and skull-meninges connections. Nat Neurosci. 2019;22:317–327.

45. Gignac PM, Kley NJ, Clarke JA, Colbert MW, Morhardt AC, Cerio D, Cost IN, Cox PG, Daza JD, Early CM, Echols MS, Henkelman RM, Herdina AN, Holliday CM, Li Z, Mahlow K, Merchant S, Muller J, Orsbon CP, Paluh DJ, Thies ML, Tsai HP and Witmer LM. Diffusible iodine-based contrast-enhanced computed tomography (diceCT): an emerging tool for rapid, high-resolution, 3-D imaging of metazoan soft tissues. J Anat. 2016;228:889–909.

46. Tun WM, Poologasundarampillai G, Bischof H, Nye G, King ONF, Basham M, Tokudome Y, Lewis RM, Johnstone ED, Brownbill P, Darrow M and Chernyavsky IL. A massively multi-scale approach to characterizing tissue architecture by synchrotron micro-CT applied to the human placenta. J R Soc Interface. 2021;18:20210140.

47. Norgaard BL, Terkelsen CJ, Mathiassen ON, Grove EL, Botker HE, Parner E, Leipsic J, Steffensen FH, Riis AH, Pedersen K, Christiansen EH, Maeng M, Krusell LR, Kristensen SD, Eftekhari A, Jakobsen L and Jensen JM. Coronary CT Angiographic and Flow Reserve-Guided Management of Patients With Stable Ischemic Heart Disease. J Am Coll Cardiol. 2018;72:2123–2134.

48. James JL, Tongpob Y, Srinivasan V, Crew RC, Bappoo N, Doyle B, Gerneke D, Clark AR and Wyrwoll CS. Three-dimensional visualisation of the feto-placental vasculature in humans and rodents. Placenta. 2021;114:8–13.

49. Celik IB, Ghia U, Roache PJ and Freitas CJ. Procedure for estimation and reporting of uncertainty due to discretization in CFD applications. J Fluid Eng-T Asme. 2008;130.

50. Roache PJ. Perspective - a Method for Uniform Reporting of Grid Refinement Studies. J Fluid Eng-T Asme. 1994;116:405–413.

51. Wang CL, Anderson C, Leone TA, Rich W, Govindaswami B and Finer NN. Resuscitation of preterm neonates by using room air or 100% oxygen. Pediatrics. 2008;121:1083–9.

52. Eckmann DM, Bowers S, Stecker M and Cheung AT. Hematocrit, volume expander, temperature, and shear rate effects on blood viscosity. Anesth Analg. 2000;91:539–45.

53. Banerjee RK, Kwon O, Vaidya VS and Back LH. Coupled oxygen transport analysis in the avascular wall of a coronary artery stenosis during angioplasty. J Biomech. 2008;41:475–9.

54. Otto JM, Montgomery HE and Richards T. Haemoglobin concentration and mass as determinants of exercise performance and of surgical outcome. Extrem Physiol Med. 2013;2:33.

55. Baidukova O, Wang Q, Chaiwaree S, Freyer D, Prapan A, Georgieva R, Zhao L and Baumler H. Antioxidative protection of haemoglobin microparticles (HbMPs) by PolyDopamine. Artif Cells Nanomed Biotechnol. 2018;46:S693–S701.

56. Tongpob Y and Wyrwoll C. Advances in imaging feto-placental vasculature: new tools to elucidate the early life origins of health and disease. J Dev Orig Health Dis. 2021;12:168–178.

57. Jackson BT, Piasecki GJ and Novy MJ. Fetal Responses to Altered Maternal Oxygenation in Rhesus-Monkey. American Journal of Physiology. 1987;252:R94–R101.

58. Carter AM. Factors Affecting Gas Transfer across the Placenta and the Oxygen-Supply to the Fetus. J Dev Physiol. 1989;12:305–322.

59. Raya JG, Dietrich O, Horng A, Weber J, Reiser MF and Glaser C. T2 measurement in articular cartilage: impact of the fitting method on accuracy and precision at low SNR. Magn Reson Med. 2010;63:181–93.

