## Supplementary Material for "An *in vitro, in utero* and *in silico* framework of oxygen diffusion in intricate vascular networks of the placenta"

#### Numerical Approach

##### Comparison to previous Numerical model

We compared our methods to Pearce et al.[1] which implements Stokes flow as well as advection-diffusion equations on COMSOL Multiphysics in three placental capillary geometries. To ensure that simulations were comparable, we used parameters used in the original study. Compared to COMSOL (finite element modelling), STAR-CCM+ uses finite volume modelling.

**Table S1: Parameters from Pearce et al.[1].**

| Parameter | Symbol | Value |
|---|---|---|
| Dynamic viscosity of blood | $\mu$ | $0.001\ Pa.s$ |
| Diffusivity of oxygen in plasma | $D_p$ | $1.7 \times 10^{-9}\ m^2/s$ |
| Diffusivity of oxygen in tissue | $D_t$ | $D_p$ |
| Oxygen advection enhancement factor | $B$ | $141$ |
| Source oxygen concentration | $c_{mat}$ | $2.24\ g/m^3$ |

##### Grid Convergence Index and Mesh Comparison

Numerical accuracy in both blood-gel domains was confirmed using a Grid Convergence Index (GCI) test [2, 3], which ensures that computed solutions are independent of the discretization approach (Table S1). Image 1 from Pearce et al.[1] was used for this study at the highest pressure drop (20 Pa). The final meshed geometry consisted of layers of prism layer cells (n = 12, stretching ratio = 1.065) at boundaries and trimmer cells elsewhere. Interface conditions were created between the flood and gel domain boundaries, surrounded by prism layer cells. Target cell sizes were 0.65 µm and 1 µm in the fluid and gel domains respectively. These settings reduced non-uniform refinement ratio GCI parameters of surface averaged WSS, oxygen transfer rate and outlet concentrations below values of 2%, which has previously been found to be adequate for mesh discretization. In general, simulations required ~5644 s of total solver CPU time to reach convergence criteria.

**Table S2: GCI parameters and results**. Data extracted from coarse, medium and fine meshes and corresponding non-uniform refinement ratio GCI values for surface averaged wall shear stress (WSS), oxygen transfer rate between the fluid and gel domains (OTR) and surface averaged oxygen concentrations on fluid outlets 1 ($c_1$) and 2 ($c_2$) respectively.

| Parameter | Cell Count | WSS (Pa) | OTR ($\times10^{-6}$ μg/s) | $c_1$ (g/m$^3$) | $c_2$ (g/m$^3$) |
|---|---|---|---|---|---|
| Coarse | 143,576 | 0.83 | 1.84 | 0.54 | 0.38 |
| Medium | 359,473 | 0.66 | 1.77 | 0.57 | 0.40 |
| Fine | 948,982 | 0.70 | 1.76 | 0.58 | 0.40 |
| GCI | N/A | 1.26% | 0.04% | 0.68% | 1.03% |

We also compared trimmer and tetrahedral core meshing models using GCI resolved mesh settings (Figure S2). For the largest inlet pressure case (20 Pa), the tetrahedral mesh overestimated the oxygen transfer rate presented by Pearce et al.[1] by 3%, while the trimmer mesh underestimated this value by 2%.

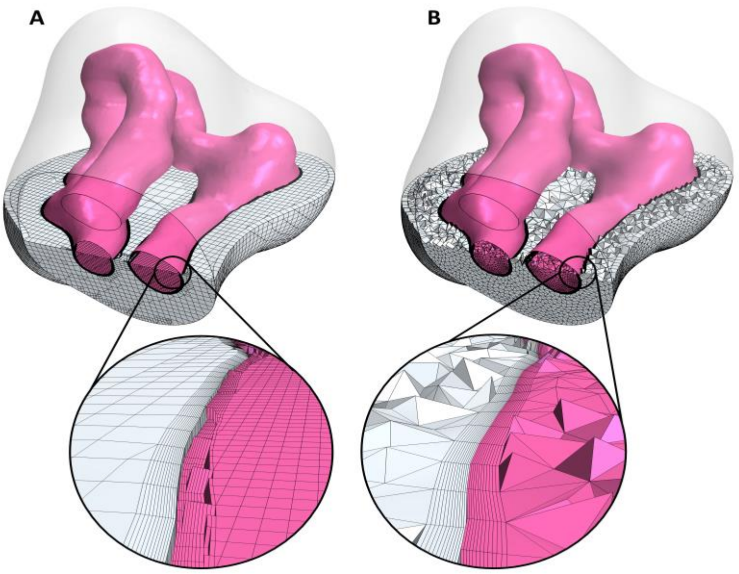


**Figure S1: Trimmer (A) and tetrahedral (B) meshes for Image 1 from** Pearce et al. [1]**.** Example zoomed cross-sections of the fluid-gel interface are shown for each mesh. Image 1 from Pearce et al.[1] were adapted under a Creative Commons licence (CC BY 4.0).

#### CALIBRATION MODEL SET-UP

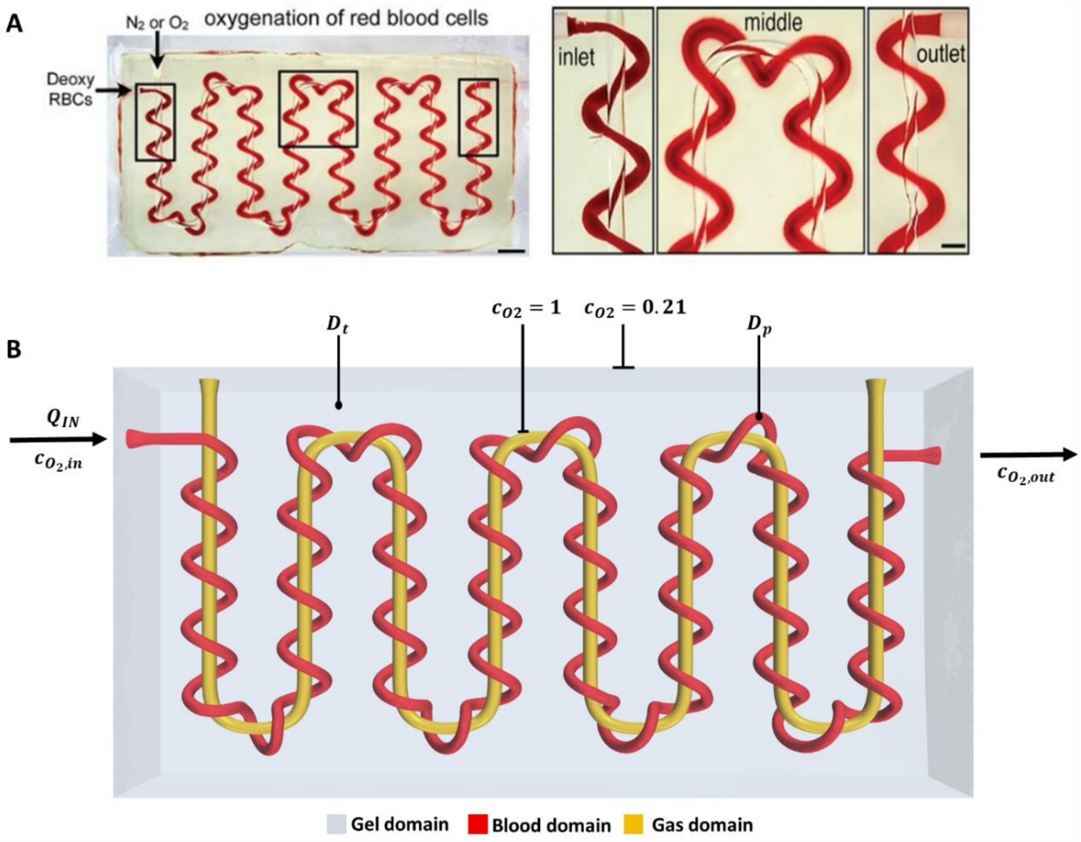


**Figure S2: Open-source serpentine-helix geometry used as the calibration model.** The size of the calibration serpentine-helix gel is 34x17x6 mm, with a serpentine channel diameter of 0.6 mm and a helix channel diameter of 0.5 mm. The inter-vessel distance is approximately 0.3 mm. The geometry consisted of gel, blood and air domains (A). For the model set-up (B), a volumetric flow rate ($Q_{IN}$) and an oxygen concentration ($c_{O_2,in}$) are assigned at the inlet, fluid and solid domain diffusivity ($D_s, D_f$) are assigned accordingly and oxygen source concentrations ($C_{O2}$) are assigned at the gas domain boundary and the external gel boundary. Mass flow is conserved and the oxygen concentration at the outlet ($c_{O_2,out}$) is calculated following solution convergence. Obtained from an open-source dataset[4] (Creative Commons Attribution Non Commercial Share Alike 4.0 International) and reproduced with permission[5] (Copyright © 2019, The American Association for the Advancement of Science).

#### MODEL SENSITIVITY

**Table S3: Sensitivity analysis of haemodynamic model parameters.** Parameters included viscosity (A), advection-enhancement parameter (B), plasma diffusivity (C) and tissue diffusivity (D). Absolute values used in primary simulations in the main paper were incrementally varied (-40%, -20%, +20% and +40%) and oxygen uptake at a high (20 Pa) and low (2 Pa) pressure drop were modelled. Absolute values of each parameter and computed oxygen gain were normalised to the baseline values used in primary simulations presented in the main paper.

**A**

| | | | Viscosity | | | | |
|---|---|---|---|---|---|---|---|
| | | Norm | 0.6 | 0.8 | 1 | 1.2 | 1.4 |
| | | Abs | 0.0042 | 0.0056 | 0.007 | 0.0084 | 0.0098 |
| Outlet O2 Conc (%) | ΔP = 2 Pa | Norm | 0.765133 | 0.89316 | 1 | 1.094431 | 1.174637 |
| | | Abs | 25.28 | 29.51 | 33.04 | 36.16 | 38.81 |
| | ΔP = 20 Pa | Norm | 0.883721 | 0.942291 | 1 | 1.055986 | 1.111111 |
| | | Abs | 10.26 | 10.94 | 11.61 | 12.26 | 12.9 |

**B**

| | | | B | | | | |
|---|---|---|---|---|---|---|---|
| | | Norm | 1 | 2 | 10 | 94 | 141 |
| | | Abs | 1 | 2 | 10 | 94 | 141 |
| Outlet O2 Conc (%) | ΔP = 2 Pa | Norm | 1 | 0.695218 | 0.350787 | 0.259383 | 0.255751 |
| | | Abs | 33.04 | 22.97 | 11.59 | 8.57 | 8.45 |
| | ΔP = 20 Pa | Norm | 1 | 0.854436 | 0.736434 | 0.710594 | 0.709733 |
| | | Abs | 11.61 | 9.92 | 8.55 | 8.25 | 8.24 |

**C**

| | | | Plasma diffusivity | | | | |
|---|---|---|---|---|---|---|---|
| | | Norm | 0.6 | 0.8 | 1 | 1.2 | 1.4 |
| | | Abs | 1.02E-09 | 1.36E-09 | 1.70E-09 | 2.04E-09 | 2.38E-09 |
| Outlet O2 Conc (%) | ΔP = 2 Pa | Norm | 0.988196 | 0.995763 | 1 | 1.004237 | 1.006961 |
| | | Abs | 32.65 | 32.9 | 33.04 | 33.18 | 33.27 |
| | ΔP = 20 Pa | Norm | 0.993971 | 0.997416 | 1 | 1.000861 | 1.002584 |
| | | Abs | 11.54 | 11.58 | 11.61 | 11.62 | 11.64 |

**D**

| | | | Tissue diffusivity | | | | |
|---|---|---|---|---|---|---|---|
| | | Norm | 0.6 | 0.8 | 1 | 1.2 | 1.4 |
| | | Abs | 3.51E-11 | 4.68E-11 | 5.85E-11 | 7.02E-11 | 8.19E-11 |
| Outlet O2 Conc (%) | ΔP = 2 Pa | Norm | 0.772397 | 0.896792 | 1 | 1.091404 | 1.166768 |
| | | Abs | 25.52 | 29.63 | 33.04 | 36.06 | 38.55 |
| | ΔP = 20 Pa | Norm | 0.887166 | 0.944014 | 1 | 1.053402 | 1.106804 |
| | | Abs | 10.3 | 10.96 | 11.61 | 12.23 | 12.85 |

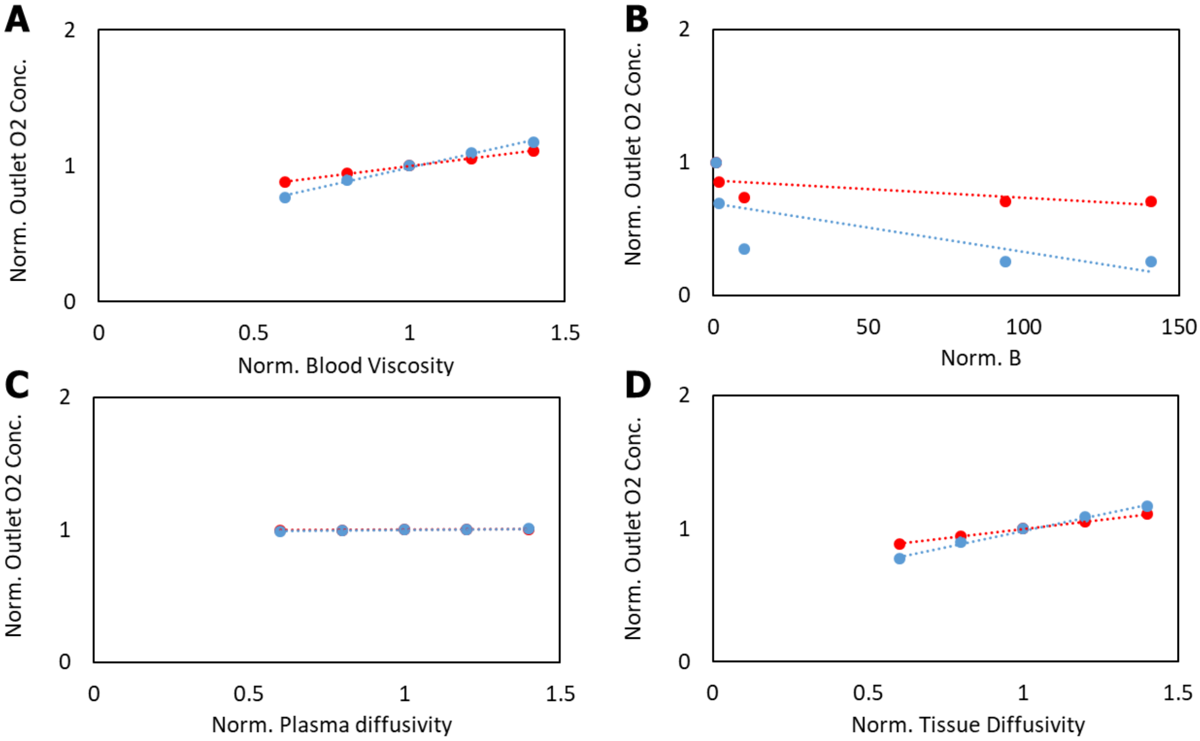


**Figure S3: Sensitivity analysis of haemodynamic model parameters.** Following normalization to baseline values of primary simulations in the main paper, the sensitivity was plotted and computed using least squares linear regressions.

**Table S4: Variation in open-source structures.** Simulations were run in 6 mathematical topologies including an axial-helix (A), a torus knot (B), a cubic lattice (C), Hilbert curves (D), a Weaire-Phelan network (E) and an alveolar network (F). Results varied between geometries, however simulations at the lower pressure drop yielded approximately 2.4 times higher oxygen gain. All geometries (A-F) were reproduced with permission[5] (Copyright © 2019, The American Association for the Advancement of Science).

| $\Delta P = 2\ Pa$ | $\Delta P = 20\ Pa$ | $\frac{c_{2Pa}}{c_{20Pa}}$ |
|---|---|---|
| A 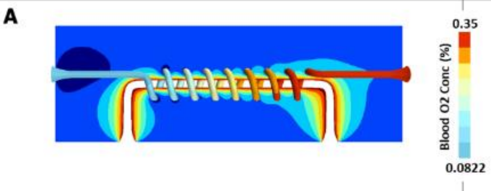  | 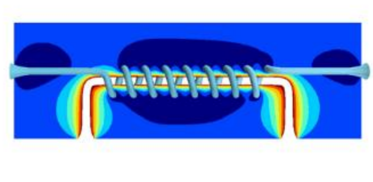 | $2.82$ |
| B 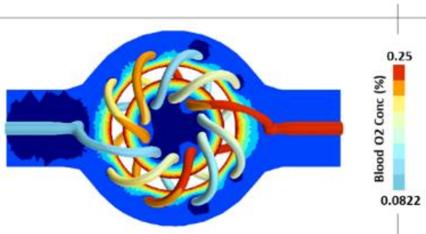  | 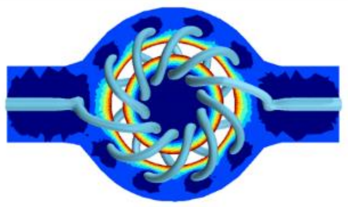 | $2.35$ |
| C 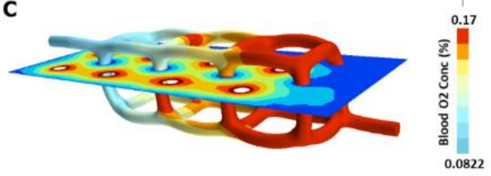  | 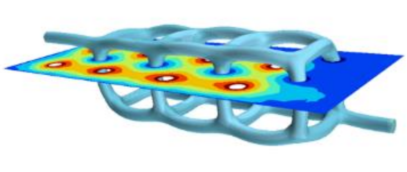 | $1.93$ |
| D 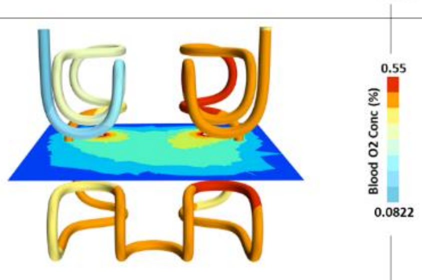  | 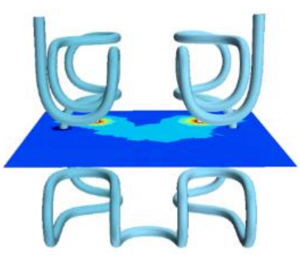 | $2.21$ |
| E 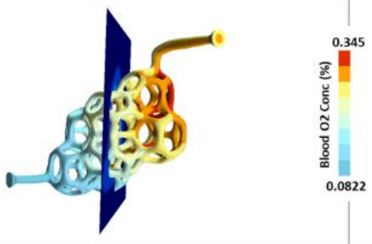  | 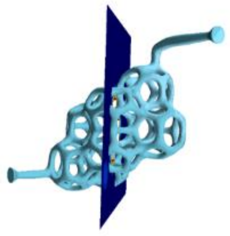 | $2.85$ |
| F 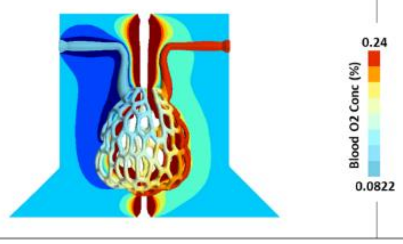  | 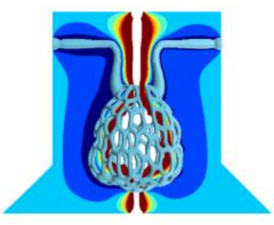 | $2.17$ |

**Table S5: Sensitivity analysis of open-source structures.** Structural parameters included gel and vessel surface area and volume, vessel centreline length, diameter, tortuosity and resistance. Simulations were run at a low and high pressure drop and oxygen gain was calculated.

| | Test Geometry | | | | | |
|---|---|---|---|---|---|---|
| Calculated Property | A | B | C | D | E | F |
| Ext Gel S.A ($mm^2$) | 1070 | 2345.49 | 938.89 | 996.71 | 512 | 781 |
| Vessel S.A ($mm^2$) | 0.0065 | 591.88 | 239.43 | 360.76 | 0.825 | 0.479 |
| Gel Volume ($mm^3$) | 1770 | 7555.11 | 1708.89 | 1761.52 | 767 | 1250 |
| Vessel Volume ($mm^3$) | 21.4 | 115.27 | 48.07 | 68.65 | 16.5 | 14.4 |
| Vessel Centreline Length ($mm$) | 72.9 | 237.9 | 53.8 | 142.4 | 28.1 | 13.1 |
| Vessel diameter ($mm$) | 0.6 | 0.79 | 0.72 | 0.7 | 0.35 | 0.26 |
| Vessel tortuosity | 0.6 | 0.94 | 0.23 | 0.69 | 0.38 | 0.57 |
| Resistance ($Pa\ m^{-3}\ s^{-1}$) | 1.58E+11 | 1.80E+11 | 2.50E+10 | 4.31E+10 | 6.85E+10 | 1.12E+11 |
| Outlet oxygen ($\Delta P = 2\ Pa$) (%) | 35.18 | 42.87 | 17.31 | 24.18 | 33.04 | 23.66 |
| Outlet oxygen ($\Delta P = 20\ Pa$) (%) | 12.47 | 19.44 | 8.96 | 10.29 | 11.61 | 10.92 |

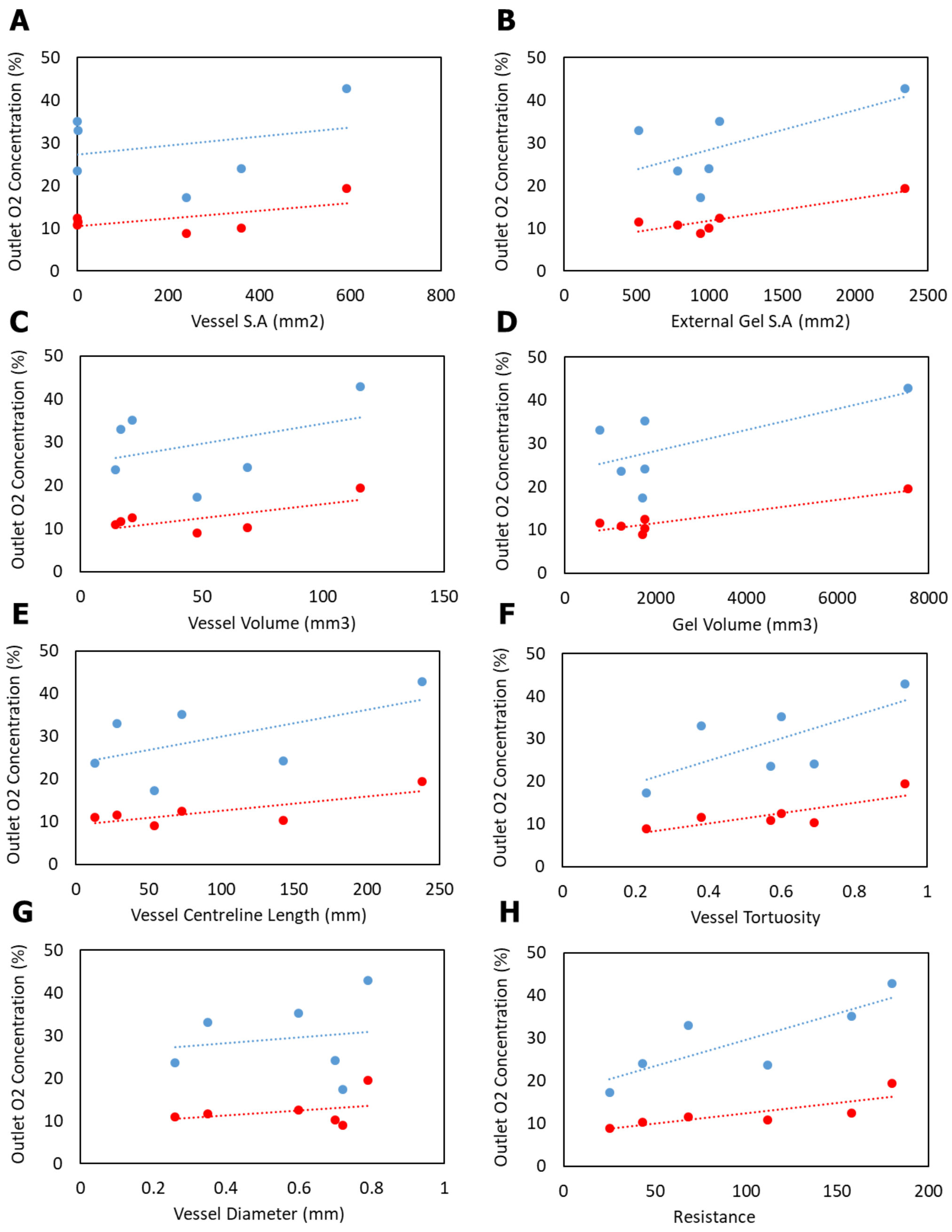


**Figure S4: Sensitivity analysis of structural parameters.** Absolute values were plotted and the sensitivity was computed using least squares linear regressions.

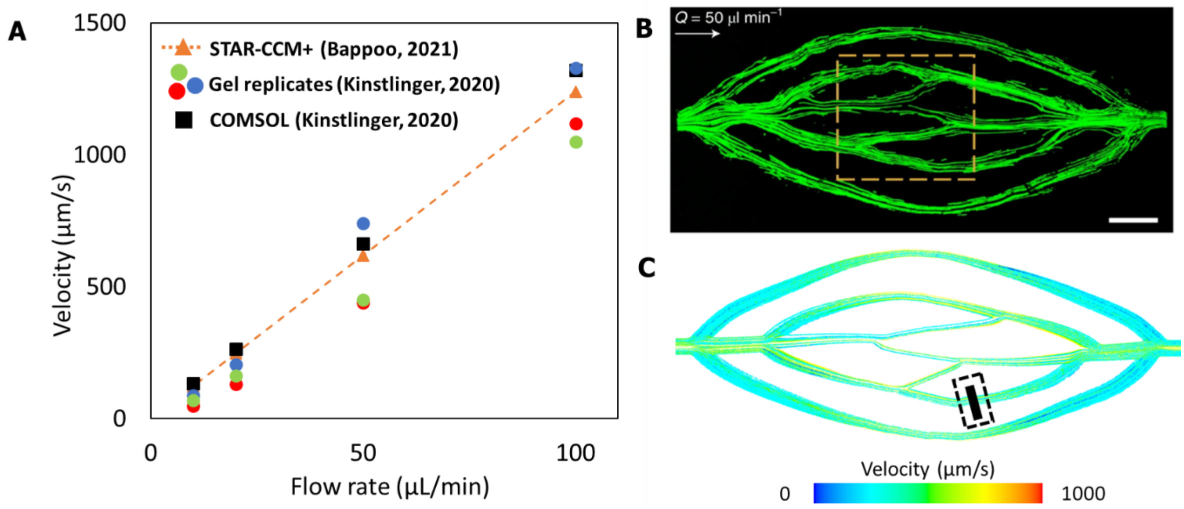


D

| Inlet flow rate (µL/min) | STAR-CCM+ | Gel average | COMSOL | % difference (Gel) | % difference (COMSOL) |
|---|---|---|---|---|---|
| 100 | 1239 | 1166.7 | 1320 | 5.8 | -6.5 |
| 50 | 619 | 543.3 | 665 | 12.2 | -7.4 |
| 20 | 248 | 166.7 | 265 | 32.8 | -6.8 |
| 10 | 124 | 70 | 135 | 43.5 | -8.9 |

**Figure S5: Experimental validation of fluid flow.** The simulation approach in this study was in close agreement to experimental data (A). PIV measurements taken in replicates of the hydrogel based model, made open source by Kinstlinger[6] (B) shown above were used for validation with the computational model (C). STAR-CCM+ simulation results were taken at the position (shown in a box in C) and compared to the PIV measurements and also to a different approach on COMSOL presented in the source paper (D). Gel replicate and COMSOL data and PIV data of idealised capillaries in B were obtained from an open-source dataset[7] (Creative Commons Attribution Non Commercial Share Alike 4.0 International) and reproduced with permission[6] (Copyright © 2020, Springer Nature Limited).

#### INVESTIGATION OF PRE-CAPILLARY OXYGEN DIFFUSION (ACROSS LABYRINTH ZONE VESSELS)

In addition to simulation configurations X and Y presented in the main text, a third scenario (Simulation Z) was investigated as it has previously been suggested that oxygenation can also occur in larger vessels such as arterioles or venules [8]. This hypothesis has not been investigated in feto-placental vessels previously as such an approach has thus been considered not physiological. We present results of this investigation.

3. **Simulation Condition Z:** Diffusion assumed though walls of feto-placental geometries (obtained from vascular casting and micro-CT) of both control (n = 1) and dex-treated (n = 1) placentas and for both normoxia (21% oxygen) and hyperoxia (100% oxygen).

Structural and haemodynamic data in rat feto-placental arterial networks were extracted at vessel diameter bins of < 50 $\mu m$, 50-100 $\mu m$, 100-150 $\mu m$, 150-200 $\mu m$, 200-300 $\mu m$, 300-400 $\mu m$ and > 400 $\mu m$. Velocities and pressures at those bins were extracted and plotted in Figure S5-A and number of vessels and relative oxygen differences (between control and dex) in Figure S5-B.

As expected, oxygen gain was pronounced in the smaller downstream vessels of both control and dex placentas (control, n = 1; dex, n = 1) both quantitatively (Figure S5-B, D), likely explained by the pressure drop and velocity deceleration of blood (Figure S5-A). This simulation variation at hyperoxia produced a -0.9% difference when compared to full domain MRI results (Table S6), demonstrating close agreement between approaches for various comparisons despite a non-physiological assumption.

**Table S6: Relative difference of control/dex differences between simulation methods and MRI measurements.** Note: Simulation Z = diffusion assumed across full feto-placental network.

| | Relative difference: Simulation Z & MRI | |
|---|---|---|
| Simulation condition | MRI full domain | MRI labyrinth zone |
| O2 at hyperoxia (100%) | -0.9 | -8.3 |
| O2 at normoxia (21%) | 12.4 | 2.5 |

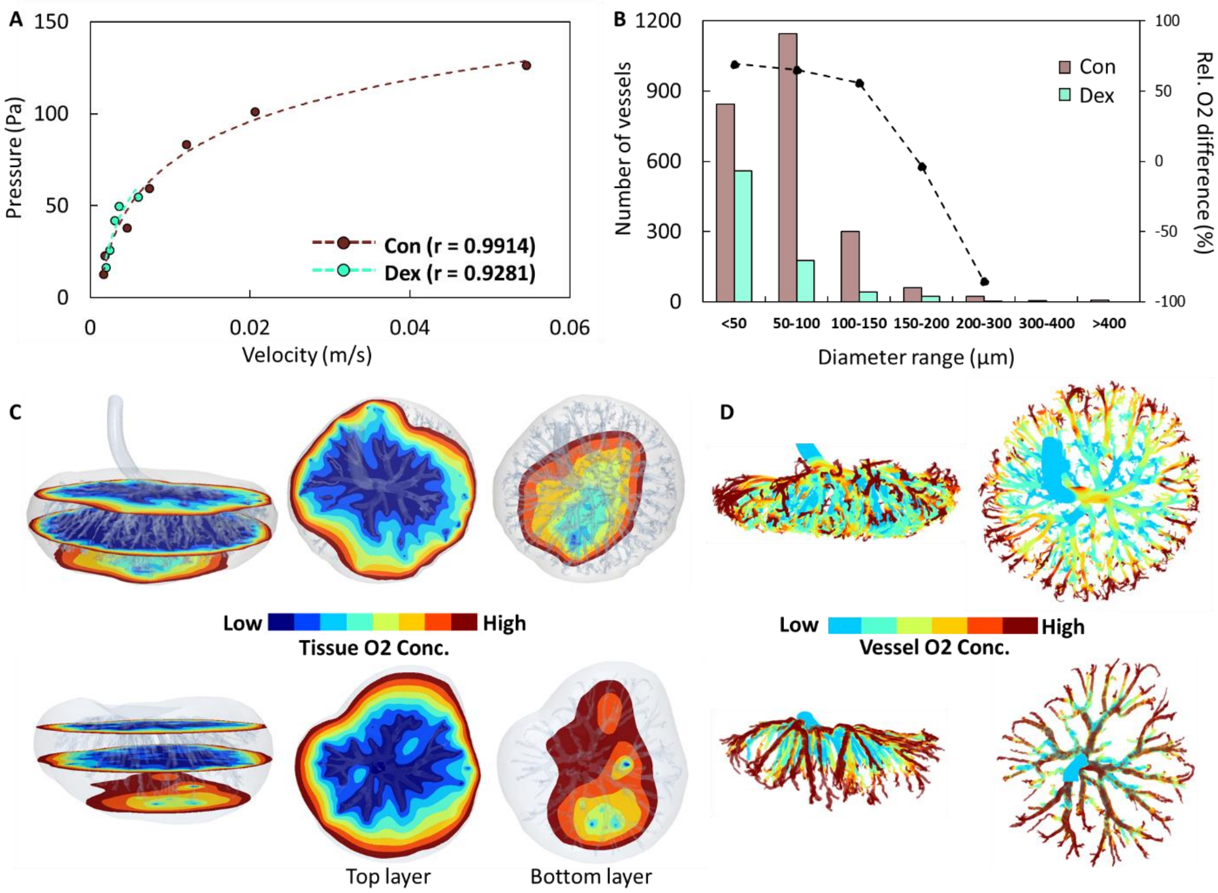


**Figure S6: Flow and diffusion across labyrinth zone vessels**. Pressure-velocity relationships at various diameters across rat feto-placental networks were fitted with a logarithmic function, showing velocity and pressure reductions across the tree structures (control, n = 1; dex, n = 1) (A). The relationship between the numbers of vessels at different diameter bins across feto-placental networks is shown along with the relative oxygen concentration difference at corresponding diameter bins (B). The control case had a lot more vessels downstream (50-100 µm range) which also coincided with a higher relative difference in oxygenation between control and dex placentas. Distributions of tissue (C) and vessel (D) oxygen concentrations are shown for control (top row) and dex (bottom row).

#### MRI Protocol

### Table S7: MRI protocols used in maternal oxygen challenge.

| Oxygen challenge | Hyperoxia via 100% medical oxygen | | | | Normoxia via medical air (21% oxygen) | | | | |
|---|---|---|---|---|---|---|---|---|---|
| **Scan no.** | 1 | 2 | 3 | 4 | | 5 | 6 | 7 | 8 |
| **Protocol** | 2D T2W anatomical imaging | 3D T2W TurboRARE anatomical imaging | T2* signal: 3D-multi-GRE sequence | T1 signal: 3D-multi-GRE sequence | | 3D T2W TurboRARE anatomical imaging | T2* signal: 3D-multi-GRE sequence | T1 signal: 3D-multi-GRE sequence | DCE-MRI with Gadovist® contrast injection (only at E21) |
| **Aim** | To locate feto-placental units | To identify fetal and placental anatomical position, and to setup field-of-view for subsequent scans | To obtain 3D-maps of the T2* relaxation maps and signals for investigating blood oxygenation | To obtain 3D-maps of the T1 relaxation maps and signals for investigating tissue perfusion oxygenation | *Change from oxygen air to medical air (3-5 min)* | To acquire anatomical position on axial and sagittal planes | To obtain 3D-maps of the T2* relaxation maps during normoxia | To obtain 3D-maps of the T1 relaxation maps during normoxia | To evaluate dynamics of placental perfusion |
| **Repetition time (TR)** | | 4000 ms | 40 ms | 20 ms | | Repeated protocol setup of scan no.2 | Repeated protocol setup of scan no.3 | Repeated protocol setup of scan no.4 | 6 ms |
| **Echo time (TE)** | | 40 ms | 1.5 ms | 1.5 ms | | | | | 2 ms |
| **Flip angle (FA)** | | 90º | 15º | 15º, 50º, 5º, 30º, 10º | | | | | 15º |
| **Echo spacing** | | 10 ms | 2 ms, 16 echoes | - | | | | | - |
| **Resolution** | | 0.25 mm x 0.25 mm | 0.5 mm x 0.5 mm x 0.5 mm | 0.5 mm x 0.5 mm x 0.5 mm | | | | | 0.5 mm x 0.5 mm x 1.5 mm |
| **Scan details** | | 2D multi-slice RARE, Fat-suppression, Respiratory trigger, One average, RARE factor = 4, Slice thickness = 1.25 mm | One average, Multiple slices covering the entire uterus were obtained, thus the number of slices varied according to the size of uterus. | One average, Multiple slices covering the entire uterus were obtained, thus the number of slices varied according to the size of uterus. | | | | | 52 cycles, Time per cycle = 20 s, One average, Field-of-view = 84 mm x 54 mm x 60 mm, Injection rate 3 mL/min |
| **Scan time** | 2 min | 5 min | 10 min | 20 min | | 5 min x 2 planes | 10 min | 20 min | 20 min |
| **Condition** | Under general anaesthesia and monitored using a PC-SAM Small Animal Monitor | | | | | | | | |

#### References


1. Pearce P, Brownbill P, Janacek J, Jirkovska M, Kubinova L, Chernyavsky IL and Jensen OE. Image-Based Modeling of Blood Flow and Oxygen Transfer in Feto-Placental Capillaries. *PLoS One*. 2016;11:e0165369.
2. Celik IB, Ghia U, Roache PJ and Freitas CJ. Procedure for estimation and reporting of uncertainty due to discretization in CFD applications. *J Fluid Eng-T Asme*. 2008;130.
3. Roache PJ. Perspective - a Method for Uniform Reporting of Grid Refinement Studies. *J Fluid Eng-T Asme*. 1994;116:405-413.
4. Miller J. Dataset for: Multivascular networks and functional intravascular topologies within biocompatible hydrogels. 2019.
5. Grigoryan B, Paulsen SJ, Corbett DC, Sazer DW, Fortin CL, Zaita AJ, Greenfield PT, Calafat NJ, Gounley JP, Ta AH, Johansson F, Randles A, Rosenkrantz JE, Louis-Rosenberg JD, Galie PA, Stevens KR and Miller JS. Multivascular networks and functional intravascular topologies within biocompatible hydrogels. *Science*. 2019;364:458-464.
6. Kinstlinger IS, Saxton SH, Calderon GA, Ruiz KV, Yalacki DR, Deme PR, Rosenkrantz JE, Louis-Rosenberg JD, Johansson F, Janson KD, Sazer DW, Panchavati SS, Bissig KD, Stevens KR and Miller JS. Generation of model tissues with dendritic vascular networks via sacrificial laser-sintered carbohydrate templates. *Nat Biomed Eng*. 2020;4:916-932.
7. Miller J. Dataset for: Generation of model tissues with dendritic vascular networks via sacrificial laser-sintered carbohydrate templates. 2020.
8. Popel AS. Theory of oxygen transport to tissue. *Crit Rev Biomed Eng*. 1989;17:257-321.
